# State-specific binding thermodynamics predicts ligand efficacy across ion-channel families

**DOI:** 10.64898/2026.09.22.753602

**Authors:** Martin Vögele, Abba E. Leffler, Kevin C. Felt, Lindsay Denluck, Edward B. Miller, Sudha Chakrapani, Lingle Wang

**Affiliations:** Schrödinger, Inc., New York, NY; Department of Pharmacology, Case Western Reserve University, Cleveland OH

**Keywords:** ligand efficacy, functional response, ion channel, binding affinity, free energy perturbation

## Abstract

Predicting ligand efficacy is a critical challenge in drug discovery, as a target’s functional response is often determined by the way a ligand shifts conformational equilibria between different functional states, a process that is particularly intricate in ion channels. We classify ligands based on the difference of their binding free energies on putative active and inactive conformations, calculated via free energy perturbation (FEP) for 78 protein-ligand pairs across six ion channels from four structural superfamilies: GluA2, GABA_A_R ρ1, α3β4 nAChR, 5-HT_3A_R, TRPML1, and KCNQ2. This approach accurately distinguishes agonists from antagonists across all these ion-channel families with large or subtle structural differences, including at membrane-facing sites, and enables quantitative prediction of maximum response and partial agonism. Importantly, we find that local binding-pocket conformations encode the bound ligand’s efficacy even when global channel states are ambiguous. Our results demonstrate that state-specific binding thermodynamics provides a robust framework for leveraging ion channel structures of diverse conformational states to elucidate mechanisms of action and to advance ion-channel drug discovery beyond simple affinity measurements, enabling the identification of new chemical matter with desired functional attributes.

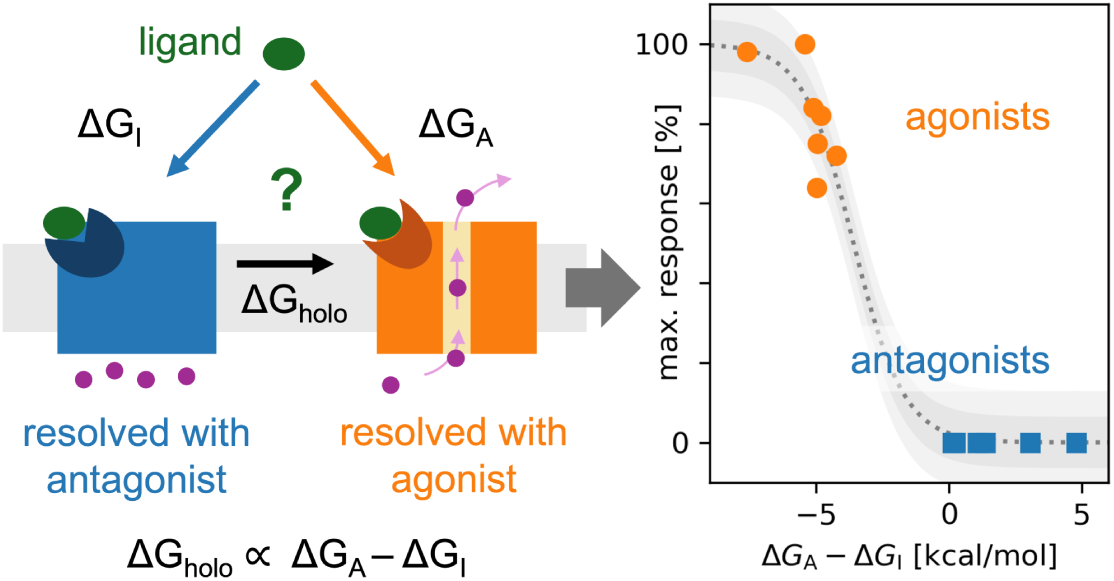

## Introduction

The function of ligands often depends not only on how strongly they bind to their target but also on how they influence the target’s conformational equilibrium. While an important initial task in early-stage drug discovery is finding molecules that bind to the target, many ligands go beyond mere binding, actively shifting the equilibrium toward specific states, with classic examples being the activation of G protein–coupled receptors (GPCRs) or ion channels. In such cases, it is essential to fully understand the mechanism of action to probe which of multiple possible target states a ligand stabilizes upon binding. With free energy perturbation (FEP) methods for estimating binding affinity now approaching experimental accuracy,^1–4^ and with structures of many classes of proteins in multiple states being routinely resolved experimentally,^5–7^ or even predicted computationally,^8–11^ a significant opportunity has emerged. These achievements open the possibility of leveraging binding FEP methods to gain more detailed and actionable insights into a ligand’s exact mechanism of action, moving beyond a simple measure of affinity and taking into account multiple target conformations.^12,13^ This can be particularly useful for virtual screening of compounds before synthesis. Predicting a ligand’s ability to influence its target’s conformational state is thus an important current challenge in computational drug design.

A systematic study across multiple targets has confirmed the relation of binding affinity and functional response and shown that we can reliably use FEP on different functional states to predict ligand efficacy.^12^ The fundamental principle connecting affinity and efficacy is that the shift in the target’s conformational equilibrium upon ligand binding is connected to the ligand’s binding affinities to the relevant states. In an idealized model with two states – we call them “active” and “inactive” here – the shift upon binding ΔΔG = ΔG_holo_ − ΔG_apo_ is thermodynamically equivalent to the difference in the ligand’s binding free energies to the active (ΔG_A_) and inactive (ΔG_I_) states, resulting in ΔΔG = ΔG_A_ − ΔG_I_ (Figure 1A/B). Based on this insight, a ligand is classified as an agonist if it binds more favorably to the active state, i.e., if ΔG_A_ < ΔG_I_. or equivalently ΔΔG < 0, and as an antagonist otherwise. In practice, the decision boundary for the classification can differ from the theoretical ΔΔG = 0, a shift that can be caused by signaling thresholds in the receptor or downstream in the experimental assay as well as by modeling artifacts in the simulations.^12,14^ We thus recommend gauging it using a few known ligands before scaling up the prediction to larger datasets. The binding free energies are calculated using Absolute Binding Free Energy Perturbation (AB-FEP). Where necessary, the protocol incorporates carefully designed restraints to maintain the receptor in its respective state. The utility of this thermodynamic rationale is also supported by similar single-target studies that have used methods based on this principle to explain or predict functional responses, including partial agonism.^15–17,14^ These achievements open the door to leveraging binding free energy methods to gain detailed and actionable insights into a ligand’s mechanism of action.

**Figure 1:**
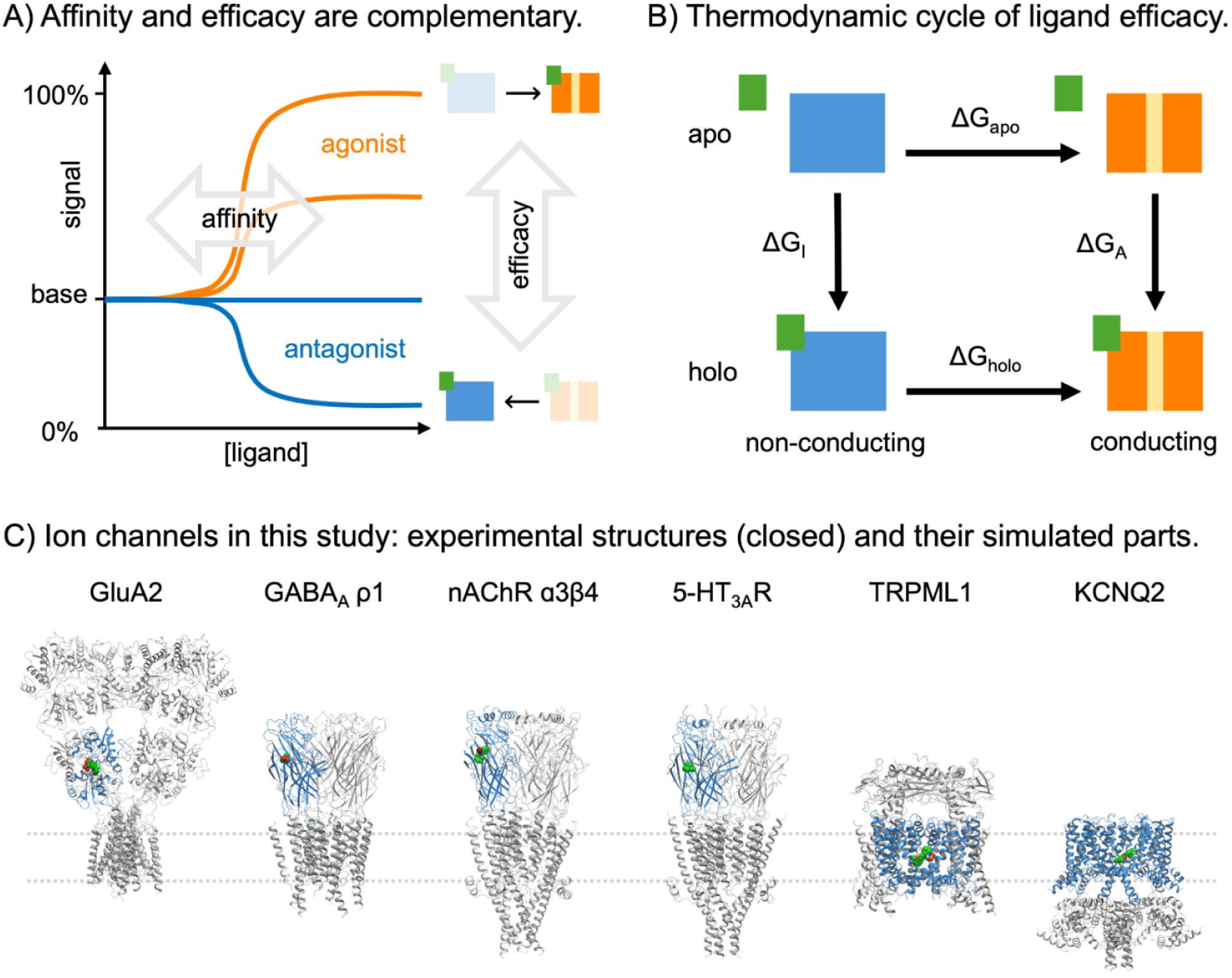
We test the possibility of predicting ligand efficacy via binding free energy to different states on six ion channels. **(A)** While affinity measures the concentration of a ligand necessary to bind to a target, efficacy determines the type (and the strength) of the effect it has on the target. Hypothetical titration curves for agonists are shown in orange and of antagonists in blue. Agonists shift the thermodynamic equilibrium of an ion channel toward the open state while antagonists shift it toward the closed state or not at all. **(B)** The thermodynamic cycle of ligand binding and channel opening, assuming an idealized two-state system. Note that the sum of free energies around the cycle is zero, resulting in *LitiG* = LiGhala - LiGapo= LiGA - LiGr **(C)** Experimental structures of the six investigated ion channels in their antagonist-bound state. The simulated sub-system is colored in blue ribbons and the co-resolved ligand is shown as spheres. The membrane is indicated with dashed lines.

While this framework has achieved high-accuracy classification across various important drug targets, including various G protein-coupled receptors (GPCRs),^12,15–17^ the retinoic acid receptor α (a nuclear receptor),^12^ and integrin αIIbβ3,^14^ an application to ion channels has not been reported due to several challenges associated with this target class. Many ion channels can exist as homo- or hetero-multimers, with binding pockets frequently located at the interface of two oligomers, and the interplay between extracellular and transmembrane domains leaves many options for functionally relevant binding sites. Different combinations of subunits in multimeric channels can exhibit different affinities, and distinct orthosteric sites can be allosterically coupled to each other, resulting in complex cooperativity of channel activation. For instance, the homopentameric serotonin 3A receptor (5-HT_3A_R) has been reported to need three of the five binding sites occupied by an agonist for activation (Hill coefficient 3.7).^18,19^ However, this number is reduced to one or two (Hill coefficient: 1.7) when adding serotonin 3B receptor (5-HT_3B_R) subunits to form 5-HT_3A/B_R heteropentamers, even though 5-HT_3B_R alone does not appear to form functional receptors.^18^ As membrane-bound proteins, ion channels are also heavily influenced by the surrounding lipid environment.^20^ Mechanosensitive channels, for example, can be directly activated by changes in curvature or tension of the lipid membrane.^21,22^ The phospholipid phosphatidylinositol 4,5-bisphosphate (PIP2) activates inward-rectifier potassium channels,^23,24^ and is a necessary cofactor for activation of several voltage-gated ion channels.^25–29^ Some channels also have binding sites for non-lipid ligands within the membrane region, exposing these to potential influence of the lipid composition.^30–33^ In general, ion channels follow a much more complex activation process than the target classes previously tackled with the functional response modeling framework introduced above. Upon agonist binding, they can undergo significant structural changes while still closed, then open for a brief time, and finally desensitize while the agonist is still bound. Even when experimental structures along this pathway are available, their assignment to one of the corresponding states is not always straightforward.^34^ Thus, the two-state model introduced above is an even stronger simplification than, e.g., for GPCRs and it is not clear *a priori* whether and how it is applicable to ion channels. Furthermore, functional data available for testing a functional response modeling protocol is scarcer for ion channels than for GPCRs. The median number of ligands per receptor reported in the GtoPdb^35^ (as of March 2026) is only 8 for ion channels versus 17 for GPCRs, and approximately 37% of known ion channel–ligand interactions are channel blockers, which do not fit the agonist-antagonist paradigm assumed by the functional response protocol. This contributes to a greater uncertainty regarding the exact mechanism of action of known ligands, as not all functional assays can distinguish between orthosteric (competitive) antagonists and non-competitive antagonists (like channel blockers), and some ligands are even thought to operate as both a competitive antagonist and a channel blocker, e.g., the nicotinic antagonist AT-1001.^36^ These inherent complexities and relative data scarcity make the study of ion channel ligand efficacy a particularly difficult undertaking.

To make this crucial yet difficult group of drug targets more amenable to ligand efficacy prediction, we demonstrate the applicability of FEP-based functional response modeling across a diverse set of six ion channels (Table 1). The selected examples are relevant targets in drug discovery and encompass binding sites in both the extracellular and transmembrane domains (Figure 1C). They also include scenarios where functional states are easily distinguishable as well as those where they show minimal difference. We briefly introduce the studied systems below and demonstrate how we can distinguish agonists and antagonists using the difference in binding affinities to different states of the respective binding pocket — somewhat simplistically designated as “active” and “inactive” — and discuss the specific challenges for each system.

**Table 1:** Experimental structures used as template structures for AB-FEP simulations. For each target we used at least one antagonist-bound and one-antagonist-bound structure while the overall state of the channel was secondary for our selection. The channel states are named as described in the articles that accompanied the resolution of the respective structure.

| Target | PDB ID | Channel State | Ligand | Effective Ligand Function |
| --- | --- | --- | --- | --- |
| GluA2 | 8SS6 | closed<sup>37</sup> | ZK | antagonist |
|  | 5NS9 | [ligand-binding domain only] | glutamate | agonist |
| GABA<sub>A</sub>R $\rho 1$ | 8OQ7 | resting<sup>38</sup> | TPMPA | antagonist |
|  | 9FRB | resting-like<sup>39</sup> | CGP36742 | antagonist |
|  | 8OP9 | likely desensitized<sup>38</sup> | GABA | agonist |
|  | 8RH7 | presumably primed<sup>40</sup> | GABA | agonist |
| $\alpha 3\beta 4$ nAChR | 6PV8 | suggested as desensitized<sup>41</sup> | AT-1001 | antagonist |
|  | 6PV7 | suggested as desensitized<sup>41</sup> | nicotine | agonist |
| 5-HT<sub>3A</sub>R | 6W1Y | non-conducting to ions<sup>42</sup> | palonosetron | antagonist |
|  | 8FSB | open-like<sup>43</sup> | serotonin | agonist |
| TRPML1 | 9HLA | closed pore<sup>44</sup> | compound 9a | antagonist |
|  | 9HJ8 | open pore<sup>44</sup> | compound 5 | agonist |
| KCNQ2 | 9IXZ | inactivated<sup>45</sup> | Ebio3 | antagonist |
|  | 9IXY | open<sup>45</sup> | Ebio2 | agonist |

## Results

### An introductory example

As an introductory example, we start with the **glutamate ionotropic receptor AMPA type subunit 2 (GluA2)**, a ligand-gated ion channel with easy-to-distinguish conformational states of the ligand binding pocket.^46,47^ While the transmembrane domain of its homo-tetramer exhibits 4-fold symmetry, its extracellular domains are organized as pairs of local dimers.^48^ Glutamate receptors are found on both neuronal and non-neuronal cells in the central nervous system where they mediate fast excitatory synaptic transmission.^46^ Antagonists for these receptors are promising candidates for treatment of epilepsy.^49^ On GluA2, we demonstrate the problem of distinguishing mechanisms of action on a dataset of competitive agonists and antagonists (that bind to the native site), and non-competitive antagonists (that bind to different sites). The challenge includes assessing both the absolute binding affinities and the difference in affinities between active and inactive states, represented by structures resolved with an agonist and with an antagonist, respectively. Given that the glutamate binding pocket of GluA2 differs significantly between these two states (Figure 2A), distinguishing between them should not be too difficult. Assuming we are looking for a competitive antagonist, we first identify ligands that principally can bind to the glutamate binding site in its inactive state and then filter out those that bind equally well or even stronger to the active state. Using only binding affinities to the inactive state (the lower value of one run with restraints and one run without), we can easily distinguish the competitive ligands from the non-competitive ones (Figure 2B). Non-competitive inhibitors, which bind to a different pocket, are mostly predicted to not bind at all to the glutamate binding pocket (ΔG > 0 or failed simulation) or to bind only very weakly. As expected for this system, agonists and competitive antagonists are easily distinguished (Figure 2C). Since the workflow uses *Absolute* Binding FEP, even ligands with structures that differ strongly from the template ligands and from each other can be correctly identified (Figure 2D). The simulations with restraints of the protein backbone even achieve the theoretically predicted threshold of ΔΔG = 0 while unrestrained simulations exhibit a small offset (SI Figure S1). Predicting accurate absolute binding free energies for this system is difficult because of the numerous potentially charged moieties present on both the ligands and within the binding pocket. Thus, in practice, the models would likely need additional refinement. Nevertheless, the model retains sufficient predictive power for functional classification due to the substantial difference between predicted binding affinities to the active and inactive states. Ultimately, this example demonstrates the general possibility to leverage binding free energy to distinguish multiple potential mechanisms of action of ion channels.

**Figure 2:**
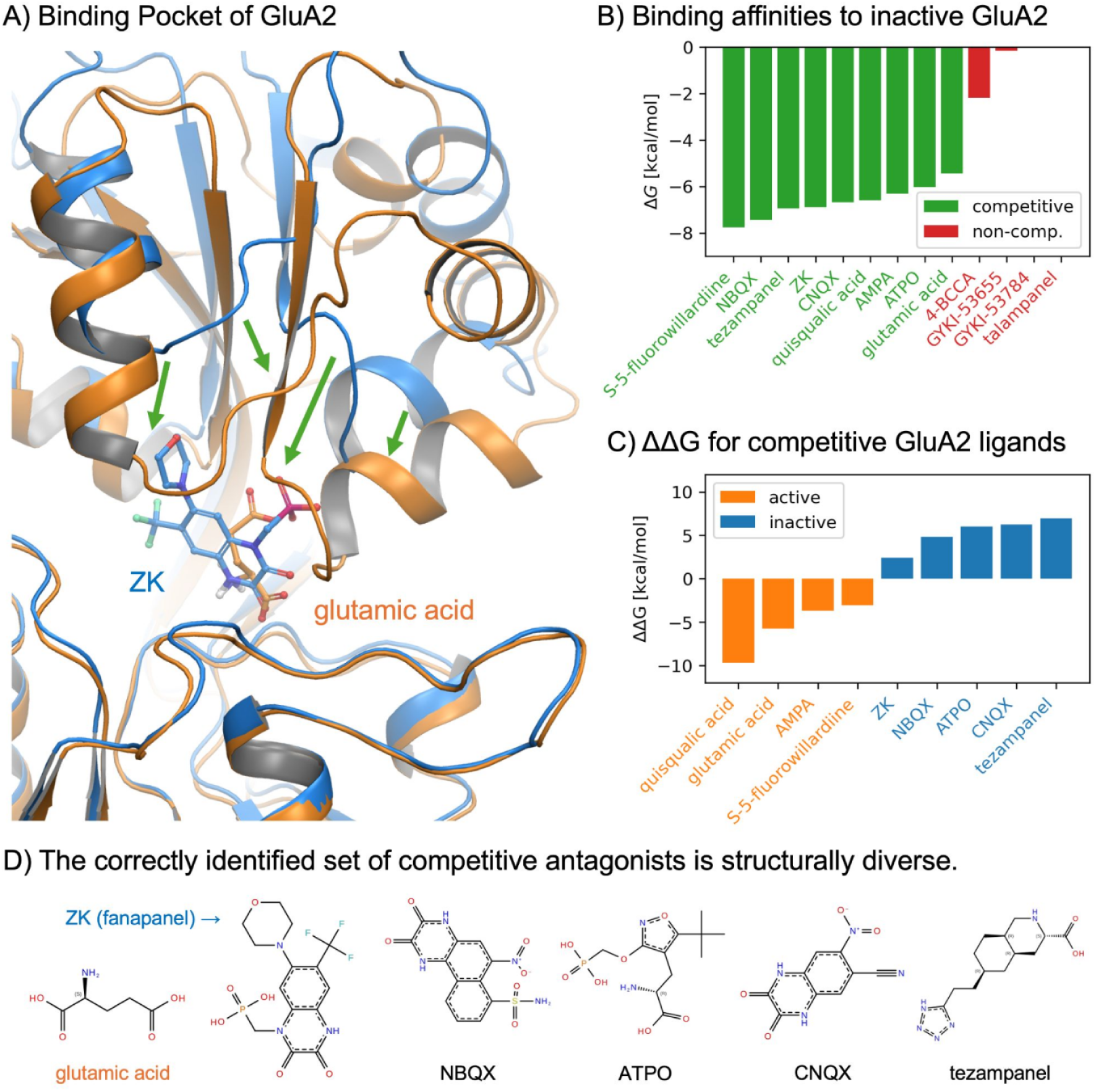
The glutamate binding pocket of GluA2 differs enough to easily distinguish agonists from antagonists among binders to this site. **(A)** Binding pocket of GluA2 bound to antagonist ZK (fanapanel) from PDB 8SS6 (blue) and bound to glutamic acid from PDB SNS9 (orange). **(B)** Predicted binding affinities to the closed, antagonist-bound state (8SS7) for all ligands (green: competitive ligands, red: non-competitive ligands). Failed simulations and positive predicted affinities are represented as O kcaljmol. **(C)** Predicted binding affinity difference between the two states for competitive GluA2 agonists (orange) and antagonists (blue). **(D)** Two-dimensional structures of the template ligands and the four other competitive antagonists in this study.

### More subtle conformational differences

We then test functional response classification for ion channels with smaller differences between the relevant states of the binding pocket. For this more difficult challenge, we choose the extracellular domains of the **gamma-aminobutyric acid type A receptor ρ1 subunit (GABA_A_R ρ1)** and the **nicotinic acetylcholine receptor subtype α3β4 (α3β4 nAChR)**. The GABA_A_R ρ1 subunit is mainly expressed in the retina where it plays an essential role in visual processing.^50,51^ Drugs acting on GABA_A_R ρ subunits (initially classified as GABAC receptors) may help in the treatment of visual, sleep and cognitive disorders.^52^ In particular, GABA_A_R ρ1 antagonists can inhibit form-deprivation myopia.^53^ The nAChRs take on numerous physiological functions in the central and peripheral nervous systems and are implicated in various neuropsychiatric disorders, including dementia and epilepsy, as well as in addictive behavior.^54^ The α3β4 nAChR in particular exhibits potential as a target for treatments of nicotine and alcohol addiction.^36,55^ Both targets are members of the Cys-loop receptor family of pentameric ligand-gated ion channels, characterized by a conserved loop structure in the N-terminal extracellular domain which is vital for ligand binding and channel gating.^56–58^ A further complication for their native binding site, beyond the small conformational differences, is that it is located at the interface of two domains. Finally, ligand binding in this pocket is driven by a cation-π interaction, an effect that depends on polarization and is not fully captured by the fixed-charge force field OPLS4.^59^ Due to these challenges, we consider this a much more difficult system for functional classification than our introductory example GluA2.

To demonstrate the influence of various subtly different receptor states, we probe four different receptor conformations of the GABA_A_R ρ1 (Figure 3A). These include two GABA-bound template structures: The first one is in a primed state, a pre-open state in which the channel has bound GABA but has not yet undergone the final structural change required to open the gate. The other structure is in a desensitized state, a post-open state in which GABA remains present but the channel’s gate is closed. The other two template structures are in a resting or resting-like state and bound to inhibitors: one to TPMPA and one to CGP36742 (Table 1). Loop C distinguishes clearly between GABA- and inhibitor-bound states. Loop B is different in the inactive, primed (pre-activated), and desensitized states, with the desensitized state being between the other two. The two inhibitor-bound states differ in the orientation of Met177 which points toward the binding site for TPMPA and away from it for CGP36742. We obtain perfect binary classification of the investigated ligands (Figure 3D) using the lower ΔG from the two inhibitor bound structures as ΔG_I_ and the lower ΔG from the two GABA-bound structures as ΔG_A_ obtained via AB-FEP using tight restraints to each structure’s backbone. Supporting a semi-quantitative agreement, the ligand whose predicted ΔΔG is closest to zero is the GABA_A_R ρ antagonist isonipecotic acid which is an agonist on the structurally similar GABA_A_R ɑ receptors.^35^ This flip of functional response upon a small receptor change suggests that the ligand’s stabilizing effect on the closed state is rather weak, which corresponds to a small ΔΔG. While ion channel activation is a complex multi-state process, the strongly simplifying two-state model still yields good predictions if the appropriate sub-states (or combinations thereof) are chosen. For our default prediction, we pool the results from primed and desensitized states together because both states are agonist-bound and structurally as well as functionally distinct from the antagonist-bound structures that represent the inactive state. A priori it is not certain which one is a better representation of the active state, although the structures already hint at the primed state because it differs more strongly from the inactive state. Breaking down the difference between primed and desensitized state, we find that indeed the primed-state structure alone yields perfect classification on its own while the desensitized-state structure alone is a slightly worse representation for the active state (SI Figure S2) Taken together, these results show the importance of probing multiple conformational sub-states and its potential to contribute to a better mechanistic understanding.

**Figure 3:**
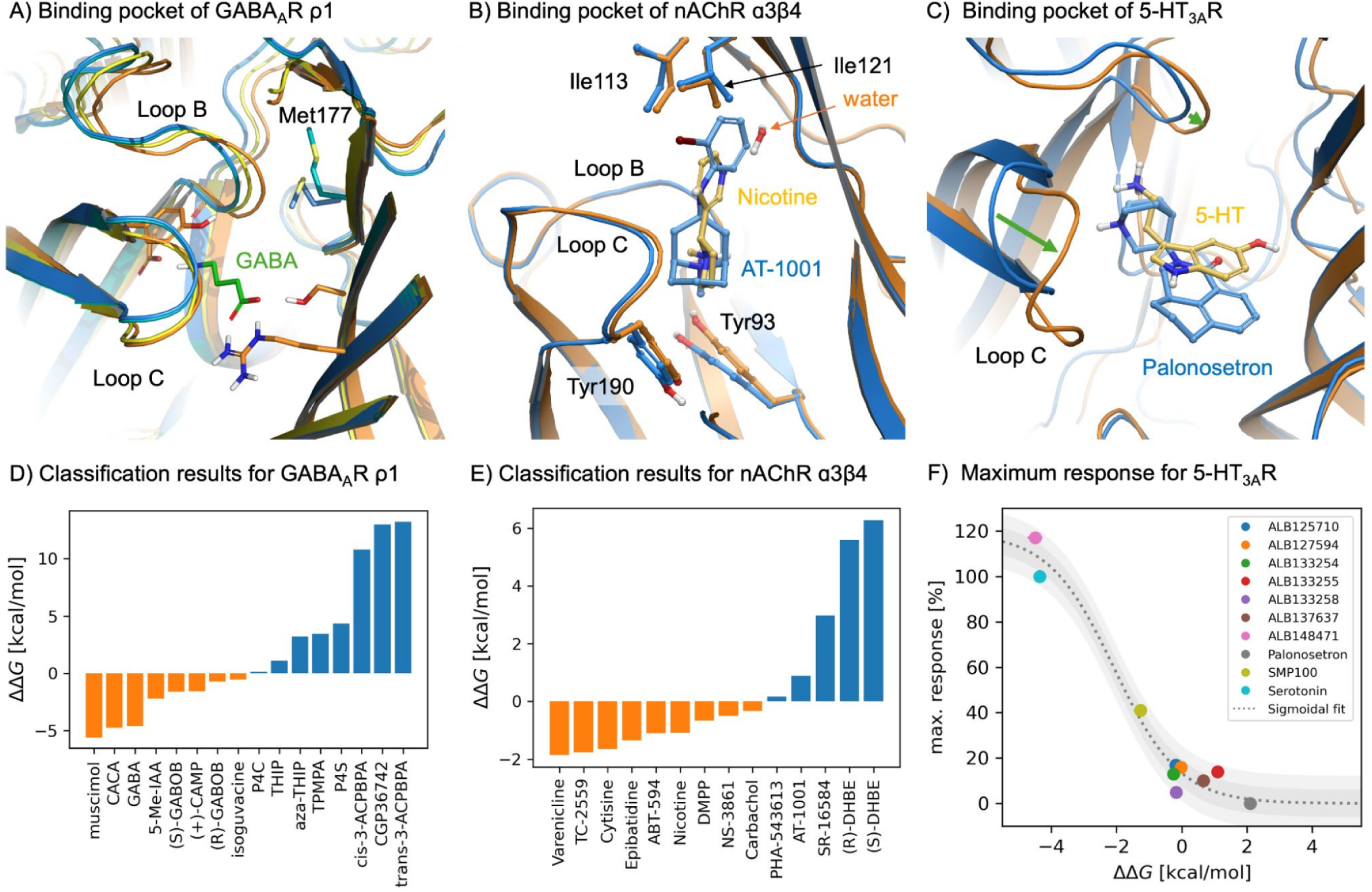
Despite only small structural differences between open and closed states, AB-FEP can distinguish agonists and antagonists for the Cys-loop receptors GABAAR pl and o.3P4 nAChR and allows for quantitative prediction of the maximum response on 5-HT3AR. **{A)** Binding pocket of the GABAAR pl in the GAB-bound primed state from PDB 8RH7 (orange), the GABA-bound desensitized state from PDB 80P9 (yellow), the TPMPA-bound resting state from PDB 80Q7 (blue), and the CGP36742-bound resting-like state from PDB 9FRB (turquoise). GABA is shown with carbons in green. **{B)** Binding pocket of o.3P4 nAChR from PDB 6PV7 (orange) with the agonist nicotine (yellow) and from PDB 6PV8 (blue) with the antagonist AT-1001 (light blue). {C) Binding pocket of 5-HT3AR in the open-like state from PDB 8FSB (orange) with agonist serotonin (5-HT, yellow) and in the non-conducting state from PDB 6Wl Y (blue) with antagonist palonosetron (light blue). **(D)** Classification results for GABAAR pl. **(E)** Classification results for the a3P4 nAChR. **{F)** Maximum response measured for various ligands as a function of the difference of binding free energy values calculated using AB-FEP, together with a fit to the theoretically expected sigmoidal relationship. The dark shaded area represents the RMSE of the fit (6% in the maximum response) and the lightly shaded area represents twice the RMSE.

We illustrate the potential to leverage even the smallest structural differences on the example of α3β4 nAChR. We use a pair of structures resolved within the same study, one with α3β4 bound to the agonist nicotine and the other bound to the functional antagonist AT-1001,^41^ which is technically a partial agonist but causes rapid desensitization.^60^ The two structures exhibit extremely small differences that are difficult to recognize by eye (Figure 3B). However, mapping out their binding sites, using Schrödinger SiteMap,^61^ shows that these small differences between the two conformations result in changes in the shape of the ligand-accessible volume in the binding pocket (SI Figure S3A–C). These changes overlap with the structural differences between the agonists (nicotine, cytisine, epibatidine) and antagonists (AT-1001, SR-16584) with the most certain poses (SI Figure S3D), suggesting that they may be used for functional differentiation. Initial tests of our AB-FEP workflow on these five ligands indeed achieve good classification using these two conformations in four repeat runs of 5 ns each (SI Figure S4). Predictions based on the average ΔG are correct for all ligands with non-overlapping standard errors, confirming a link between subtle binding pocket structure differences and ligand function. However, overlapping standard deviations suggest that a single 5 ns run may not suffice for accurate prediction. For our final prediction on the entire data set, we thus use a 10 ns run instead. We then achieve perfect classification on the full dataset using only the difference of ΔG between these two states (Figure 3E). This also holds for ligands with scaffolds different from the co-resolved ligands and, remarkably, even those with side groups that clearly reach beyond the core binding pocket: PHA-543613 and (S)/(R)-DHBE. As a notable semi-quantitative agreement, PHA-543613 is predicted to be near zero, which is consistent with its behavior as a very weak antagonist on ganglion-like nAChRs, where an inhibition of only 13% has been experimentally observed.^62^ This all-around successful classification shows that even small differences in the template structures can be used to distinguish ligands with different functional responses, provided that these differences are relevant for the investigated effects.

Successful classification of agonist versus antagonist requires accurate calculation of the binding free energies of each ligand to the respective receptor conformations. Considering the challenges associated with the absolute binding free energy calculations, the requirement that the binding free energies to both receptor conformations have to be accurate, and the fact that some ligands only slightly favor the active or inactive state, it is remarkable that all the ligands for the the α3β4 nAChR are correctly classified even for these few with calculated ΔΔG close to zero. Since it requires two receptor conformations and the classification ΔΔG threshold is close to zero, the α3β4 nAChR is an ideal system to analyze the errors of the calculations, and how likely the ligands are correctly classified based on the errors of the calculations. To connect the errors in the free energy calculations with the likelihood a ligand is correctly classified as an agonist or antagonist, we built a statistical model that separates the error of an AB-FEP calculation into statistical sampling error (noises in repeated calculations) and systematic error (deviation between experimental and average of repeated calculations), and quantifies how the sampling error and the correlation between the systematic errors associated with the two receptor conformations affect the likelihood of correct classification (SI Text). As an upper bound for the sampling error of ΔG_A/I_ we use 0.325 kcal/mol, estimated from the median standard deviation in the short repeat runs that we scaled by a factor of 1/√2 to account for the longer simulation time (SI Figure S4B). As a lower bound we use the 0.1–0.2 kcal/mol Bennett error obtained from individual AB-FEP runs (SI Figure S4C). With those levels of sampling errors and assuming there is no systematic error, we would expect the perfect classification with a probability of 40% using repeat-run errors and 86% using Bennett errors, and at most one wrong prediction with 80% to 100% probability, respectively. Clearly, the error between a converged AB-FEP result and experimental value (systematic error) is not 0, and a systematic error of ∼1 kcal/mol for the individual ΔG values is expected for an AB-FEP result based on earlier studies (SI Figure S4D).^3^ The systematic error is caused by inaccuracies in the ligand pose, protonation states, and force-field parameters as well as insufficient sampling of potential conformational fluctuations of the protein. If the systematic errors in ΔG_A/I_ on the two receptor conformations propagated to ΔΔG independently the same way as the sampling uncertainty, it would correspond to a much lower likelihood of success: 10% for perfect classification and 36% for not more than one wrong prediction. However, since the systematic errors in ligand or protein protonation states and force-field parameters are largely the same in the two simulations for the same ligand on the two receptor conformations, we expect a strong correlation in the systematic errors in the two ΔGs from which the ΔΔG is calculated. Based on the perfect classification performance observed for α3β4 nAChR, our model suggests a high correlation between the systematic errors in the two AB-FEP ΔGs for each ligand on the two receptor conformations, e.g., perfect classification becomes more likely than not for a correlation factor of 0.94 with sampling errors coming from the Bennett errors. This makes sense as the protonation states and force field parameters in both α3β4 nAChR simulation setups are exactly the same and the starting poses of the ligands only differ minimally. Any errors associated with these parameters are of systematic nature and thus largely the same in both simulations, canceling each other out in the difference ΔΔG = ΔG_A_ – ΔG_I_.

We go beyond binary classification and investigate agonism quantitatively for the **serotonin 3A receptor (5-HT_3A_R)**, another Cys-loop receptor. As the only class of serotonin receptors that are ion channels instead of GPCRs,^63^ 5-HT_3_ receptors play a crucial role in both the central and peripheral nervous systems where they mediate fast excitatory neurotransmission, implicating them in numerous physiological processes like the regulation of anxiety, nausea, and vomiting.^64,65^ Recently published data of mouse and human 5-HT_3A_R ligand efficacy contains consistent measurements of the maximum functional response (E_max_),^43,66^ ensuring that observed differences in ligand efficacy are primarily attributable to the intrinsic properties of the ligands and the channel-ligand interactions rather than variability of experimental conditions. As templates to probe the ligands against, we use the two most extreme conformations identified on the resolved conformational spectrum of m5-HT_3A_R: a non-conducting state in complex with antagonist palonosetron and an open-like state in complex with serotonin.^42,43^ From the theoretical two-state model underlying our workflow,^12^ we expect a sigmoidal relationship between the functional response and the free energy difference (see Methods). Indeed, the measured E_max_ agrees well with a fitted sigmoidal relationship when plotted against ΔΔG determined from the FEP predictions for the ligand’s affinity for these two extreme conformational states (Figure 3F). We can thus separate out the partial agonist SMP100 from the full agonists and a group of antagonists and very weak partial agonists. Due to the sigmoidal relationship, the transition zone between minimal and maximal signal is inherently narrow,^3^ which demonstrates the general difficulty in the prediction of precise E_max_ values in this critical region. From the trajectories of the FEP simulations, we can gain mechanistic insights by comparing the interactions with the binding pocket of the congeneric polycyclic tertiary-amine ligands between the two states. We identify Asp202 as a crucial residue for agonism (SI Figure S6A), consistent with previous findings.^43^ Ligands that contain a pyrrole or pyrazole ring with a hydrogen atom at N4 can form a hydrogen bond with Asp202, stabilizing the inward motion of Loop C (SI Figure S6B). The exception is ALB137637, an enantiomer of SMP100. Due to the different conformation of stereocenters (SRS instead of RSR), the charged amine anchors ALB137637 differently in the pocket such that the resulting tilt allows a water bridge from N3 to Asp177 (SI Figure S6B). The free energy reduced by the hydrogen bond to Asp202 in the open-like state is at least partially offset by breaking this water bridge, which can explain this ligand’s lack of agonism despite the ability to form the Asp202 hydrogen bond. This serves as a reminder that even if we think we have identified a pattern of interactions that explain agonism, outliers to this can appear when other parts of the ligand change the balance of interactions. FEP can identify these outliers because it takes all relevant interactions into account. For this receptor we have to caveat that the maximum response and binding affinity were measured on a human receptor while the structures used from FEP were from a mouse. This might have contributed to a shift in absolute values of the binding free energy (SI Figure S7A) or to the shift of the sigmoidal curve along ΔΔG, although also reorganization free energy and force field inaccuracies can contribute to these. The two mutated residues with the most likely effect on agonism (Ile201 and Asp202) are conservative mutations (SI Figure S7A–C). The decisive hydrogen bond can be formed as well with glutamic acid Glu224 as with Asp202, which explains why the sigmoidal distribution of the maximum response is preserved for the ligand series investigated here. This may not hold for even bulkier ligands, such as the recently resolved vortioxetine.^67^ Thus, in practice, we recommend using structures from the same species if possible. Still, these results demonstrate that our workflow can even provide quantitative insights into the degree and the mechanism of a ligand’s agonism.

### Membrane-facing binding sites as a particular challenge

Binding pockets in the transmembrane domain that face the lipid membrane present a distinct and challenging problem, as exemplified by the **mucolipin TRP cation channel 1 (TRPML1)** and of the **potassium voltage-gated channel subfamily Q member 2 (KCNQ2)**. TRPML1 (also known as MCOLN1) is a non-selective cation channel found primarily in endosomal and lysosomal membranes where it plays a crucial role in membrane trafficking, signal transduction, and ionic homeostasis.^68^ It mediates autophagy events involved in the pathogenesis of cancer and myocardial ischemia reperfusion injury, which makes it a potential target for the development of therapeutics against these diseases.^69^ KCNQ2 (also known as Kv7.2), a subunit of the KCNQ family of voltage-gated potassium channels, is expressed mainly in the central and peripheral nervous systems. Together with other KCNQ isoforms, it is vital for regulating neuronal excitability via the M-current (a slow, non-inactivating potassium current).^70^ Due to the association of KCNQ2 mutations with various forms of epilepsy, it is a significant drug target for neurological disorders.^71,72^ A particular challenge for studying ligand activity on these channels is the membrane-facing location of their pore-adjacent hydrophobic cavities.^30–33^ Interactions of the ligand with lipids complicate binding affinity calculations because they converge more slowly than both the stable and generally well-defined interactions in conventional protein binding pockets and the quickly fluctuating ligand-water interactions on aqueous surfaces. While classification of ligands for TRPML1 was found to work well without significant complications, the more challenging example of KCNQ2 serves as a valuable case study to illustrate the boundaries, weaknesses, and necessary prerequisites for the FEP-based functional response workflow.

Sufficient experimental data on channel opening and binding poses enable a thorough evaluation that shows accurate classification of ligands that bind to the membrane-facing fenestration pocket of TRPML1. For this target, we benefit from a consistent ligand efficacy dataset and poses from resolved structures, avoiding possible uncertainty in ligand poses from docking.^44^ In contrast to the systems with binding pockets in the extracellular domain discussed above, systems with membrane-facing binding pockets are set up using a lipid membrane made of 1-palmitoyl-2-oleoyl-sn-glycero-3-phosphocholine (POPC). Due to the uncertainty inherent in the highly variable lipid positions, we tested the impact of the co-resolved lipid. We ran two types of TRPML1 simulations for each ligand and state: once with the co-resolved lipid in the experimental structure and once with the POPC membrane modeled around the respective resulting system. The simulations without the co-resolved lipid provide another opportunity for the modeled-in lipids to settle into favorable positions around ligands that do not match the co-resolved lipid well. We found that while individual runs were not always robust, the failures only occurred in the ligand’s unfavored state due to clashes between the ligand and the incompatible binding pocket, i.e., in the active state for an antagonist or the inactive state for an agonist, thus preserving a correct separation of agonists and antagonists in both cases (SI Figure S8). Using the lowest ΔG per ligand and state from all successful simulation runs yielded excellent overall classification and even a good quantitative agreement with experiments (Figure 4C). A detailed investigation between the maximal percentage of response and the magnitude of ΔΔG is possible because all studied ligands are from a single, consistently measured electrophysiology study including maximum response values for agonists.^44^ Ordering the ligands along ΔΔG separates the two strongest agonists (compounds 2 and 7) from a group of somewhat weaker agonists, from the low-potency compounds (1b, 4b, and 9b), and from the antagonists. A sigmoidal fit assuming E_max_ = 0 for antagonists and excluding the low-potency compounds yields an RMSE of 6.5%, similar to 5-HT_3A_R. Although no binding affinity measurements are available for this dataset, we can use the potency measured as IC_50_ for antagonists and as EC_50_ for agonists as a proxy. We find a roughly linear correlation between –pIC_50_ and –pEC_50_, respectively, and the lowest ΔG. The linear fit with an R^2^ of 0.68, RMSE of 1.69 kcal/mol shows a predictive power worse than the expected 1 kcal/mol which is not surprising in the difficult membrane environment and given that potency and binding affinity are not exactly correlated. Despite this uncertainty in the absolute affinity, the functional response is modeled very accurately in comparison, again similar to the results for 5-HT_3A_R. Offering both reliable overall classification and quantitative accuracy, this study shows that with sufficient structural knowledge, FEP-based functional response prediction can excel even in the challenging environment of a lipid membrane.

**Figure 4:**
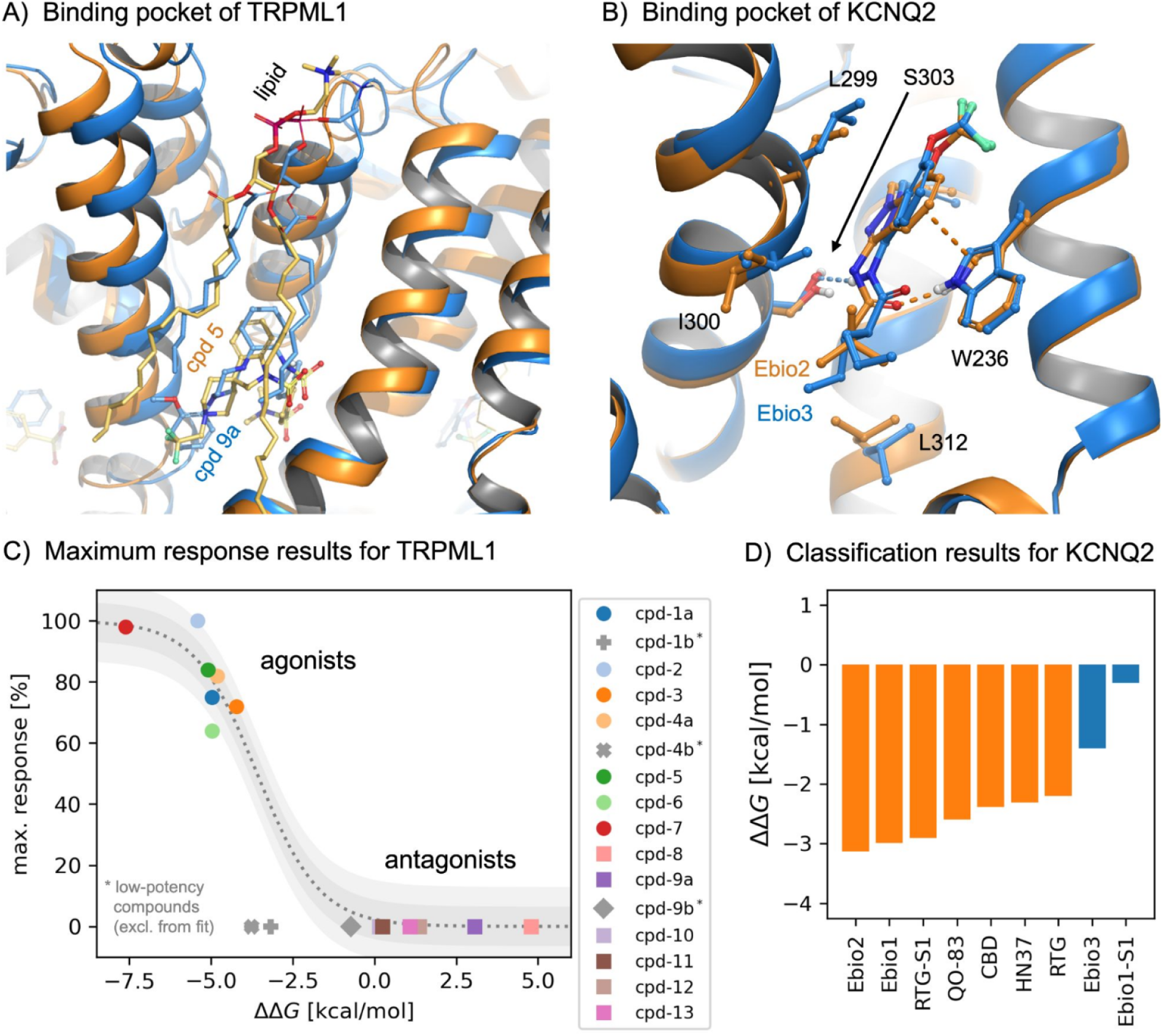
Classification for ligands targeting the pore-adjacent hydrophobic cavities of the ligand-gated ion channel TRPML1 and the analogous pocket of voltage-gated ion channel KCNQ2 succeeds qualitatively when the threshold is adjusted for each target. **(A)** Binding pocket of TRPML1 in the open state from PDB 9HJ8 (orange) with agonist compound 5 and a co-resolved lipid 3-sn-phosphatidylcholine (yellow) and in the closed state from PDB 9HLA (blue) with antagonist compound 9a and the same co-resolved lipid (light blue). **(B)** Binding pocket of KCNQ2 in the open state from PDB 9IXY with agonist Ebio2 (orange) and in the inactivated state from PDB 9IXZ with antagonist Ebio3 (blue). **(C)** Maximum response on TRPML1, measured for various ligands as a function of the difference of binding free energy values calculated using AB-FEP, together with a fit to the theoretically expected sigmoidal relationship. The dark shaded area represents the RMSE of the fit (6% in the maximum response) and the lightly shaded area represents twice the RMSE. **(D)** Classification results for KCNQ2.

Applying a similar protocol to the pore-domain binding site of KCNQ2 separates agonists from antagonists but the classification threshold deviates from the theoretical ΔΔG = 0. We start from two structures that were consistently resolved within a single study, one with the agonist Ebio2 (9IXY) and one with the antagonist Ebio3 (9IXZ).^45^ While the difference in binding pocket between them is small, we leverage it to predict the functional response (Figure 4B). Comparison to KCNQ2 structures resolved with other ligands, e.g., with the agonist CBD (PDB 8J01)^31^ and the antagonist Ebio-S1 (PDB 8X43),^32^ showed a range of possible conformations within the open and closed states, prompting us to choose flat-bottom harmonic restraints (see Methods). Since no lipids were co-resolved in the template structures, we obtain six different lipid configurations for each of the two states from independently equilibrated membranes (see Methods). To probe which configuration is most likely to accommodate each ligand best, we run AB-FEP for each ligand with all six configurations and choose the lowest ΔG to represent the corresponding state. The resulting values of ΔΔG are all negative but agonists are well separated from antagonists (Figure 4D). With a classification threshold of ΔΔG ≈ −2 kcal/mol instead of the theoretical ΔΔG = 0, we achieved correct classification. In additional simulations with narrower restraints, we find that the structural landscape of KCNQ2 reaches beyond just the two template structures and restricting it too strongly can lead to a degradation in performance (SI Figure S10), justifying our choice of flat-bottom restraints. Note that both CBD and HN37 can bind in pairs, with the second identical molecules occupying peripheral parts of the binding pocket.^31^ Because we probed only one molecule for the classification, our result neglects the effect of the second one. Additional AB-FEP simulations of the second molecule in presence of the first one for each of these two ligands showed no strong difference between the two states (SI Figure S11). Taking them into account would not change the classification, confirming that agonism is mediated by the first molecule in coordination with PIP_2_. However, our workflow cannot capture the antagonistic effect of the second HN37 because it acts by displacing the essential cofactor PIP_2_ and not by directly shifting the conformational ensemble of the receptor. Overall, the results for KCNQ2 show how even in difficult systems that do not follow the ideal two-state model with separation at zero ΔΔG perfectly, we can achieve a practically useful separation of ligands by their functional response.

## Discussion

We adapted a previously established FEP-based functional response modeling workflow to ion channels, a particularly challenging target class due to their multimeric architecture, their complex activation and desensitization cycle, and the variety of mechanisms by which ligands can modulate their activity. To achieve this goal, we applied various modifications to model ligand efficacy for binding pockets in both the extracellular domain and the more complex transmembrane domain, specifically the pore domain. The successful modeling of various scenarios shows that the main rationale of functional response modeling is in principle applicable to any system where ligands function via altering the conformational equilibrium of the target protein. Here we discuss the lessons learned from the overall performance and from each type of target as well as the limitations of the workflow.

As a general principle, the ligand binding pocket itself turns out to be the primary region of interest for the prediction of functional response. Several of the ion channels investigated here were resolved with the channel’s pore in functionally identical conformational states, the agonist-bound structure of GluA2 even without the pore domain at all. Yet, they bind functionally disparate ligands (agonists versus antagonists, Table 1). In these instances, the structural differences are highly localized within the binding pocket itself. Through the application of specific restraints in our FEP simulations, these localized differences are effectively propagated to the probed ligands, allowing for a precise differentiation of their functional effects. The question about whether a binary ligand classification can be mapped well on a two-state model (with sub-states if appropriate) is thus not decided by the conformation of the pore in the template structures but by the conformational landscape of the ligand binding pocket – which in some cases can be effectively decoupled. This finding highlights that even when global structural changes are not evident in the template structures, their conformational signature within the binding site remains a robust predictor of efficacy.

Notably, predicting a ligand’s preference for a specific receptor state is more accurate than standard comparisons of FEP to binding affinity measurements would suggest, because systematic sources of error correlate between receptor states and in many cases cancel out at least partly. Across all targets, we can separate agonists from antagonists even when the gap between them is significantly smaller than 1 kcal/mol. This becomes especially clear for the receptors with only small geometric differences between the binding pocket, particularly ɑ3β4 nAChR, where we obtained a result that suggests sampling noise to be the major source of uncertainty while systematic errors are strongly correlated (SI Figure S5). In these systems the setup of the two states is so similar and the differences so small that many of the systematic errors that account for the uncertainty of the absolute binding free energy fortuitously cancel each other out when calculating ΔΔG. Targets with larger conformational differences between states, e.g., GluA2, are more likely to require different setups (protonation states, cofactors, etc.) but the larger difference also makes the states easier to distinguish. In some cases, they do not cancel each other out completely but the remaining artifact is the same for all ligands, or at least a large subset. Then it contributes to a shift of the classification threshold away from the theoretical zero. This balance between more easily distinguishable differences and more strongly correlated errors accounts for the good performance observed for our diverse set of targets and inspires optimism that we can apply the functional response modeling principles to an even wider array of target classes.

The GluA2 example serves as a primary illustration of the general principle for probing multiple potential mechanisms of ligand action by leveraging binding free energy differences. Taking into account multiple potential binding sites and conformational states in computational predictions enables researchers to design for specific outcomes; for example, finding a competitive antagonist involves identifying ligands that bind to the inactive state of the native binding site and then filtering out those that bind more favorably to the active state. Non-competitive ligands are distinguished by their predicted weak or failed binding to the glutamate pocket. In the next step, competitive ligands (agonists and antagonists) are separated based on the difference in their binding free energies between the active and inactive states. This distinction between binding affinity prediction and functional response prediction is analogous to the problem that a functional assay alone can usually not distinguish competitive from non-competitive binders while binding assays, even when they are site-specific, do not necessarily show the functional response of the target. Free energy calculations can also be used to explain experimental results in the absence of one of these two complementary assay types. The appeal of our workflow here is that it only requires additional runs of the same software and not the setup of a completely different protocol. It is important to note, however, that the use of restraints to stabilize the receptor in a specific state, while often necessary for functional classification, can potentially distort the absolute value of the binding free energy. Probing for affinity and for efficacy may thus occasionally require different settings. In summary, this example demonstrates how differences in predicted binding free energy to various receptor states can be used analogous to differences in binding free energy to different binding sites to distinguish between different mechanisms of ligand action.

The small structural differences between the agonist-bound and antagonist-bound states of the extracellular domain of Cys-loop receptors present a unique challenge as they necessitate the use of strong restraints for effective distinction. This approach slightly differs from the one used earlier for GPCRs, which employed flat-bottom restraints derived from MD simulations.^12^ An advantage of using strong restraints is that it allows us to focus only on a part of the system, in this case two adjacent extracellular domains. This approach significantly reduces the computational burden, which is particularly useful for large multimeric ion channels. It also eliminates the need for computationally expensive, long MD simulations of the entire multimer in a membrane which due to their size and complexity are difficult to set up and take long to converge. However, a drawback is the potential to overlook some conformational sub-states, especially when the experimental coverage of the conformational landscape is limited. In return, when multiple sub-states are known, strong restraints allow us to probe them separately which can contribute to a better understanding of their respective function, as demonstrated for the primed and desensitized states of the GABA_A_R ρ1. The finding that the primed state offers a superior representation of the active state enhances our understanding of the ion channel activation cycle. The part of the binding pocket that is shifted slightly toward an inactive conformation (Loop B) might be part of the allosteric pathway that connects the ligand to the desensitization gate which is located on the opposite side of the membrane.^58^ Ultimately, this example illustrates how FEP can be leveraged beyond efficacy prediction to test structural hypotheses regarding the underlying mechanisms and how more than one structure per functional state can be probed.

An ideal use case for strong restraints are the extremely subtle differences between the agonist-bound and antagonist-bound states of the nAChR, which make functional response prediction in the design process particularly difficult. In fact, such small differences — given that they accurately represent the activation mechanism — can simplify the investigation by reducing the need to sample multiple sub-states. We caveat for the α3β4 nAChR in particular that the known conformational landscape may only be adequate for the specific types of ligands studied. This is because the two co-resolved ligands are technically both agonists whose functional difference stems from the rate at which they induce desensitization.^41,60^ Data on this kind of small-molecule α3β4 nAChR antagonists is limited and the only probed antagonists that reach beyond the core binding pocket are PHA-543613 and (S)/(R)-DHBE. Peptidic nAChR antagonists, α-conotoxins, stabilize a binding pocket conformation that differs more strongly from the nicotine- and AT-1001-bound conformations, as suggested by mutagenesis and by structures of the α3β4-mimetic acetylcholine-binding protein, the α7 nAChR, and the muscle-type nAChR.^73–75^ The structural model would have to be extended by the structure of such a state in order to better generalize to this type of peptidic ligands. To conclude, strong restraints are a necessary and practical approach for studying systems with subtly different functionally distinct states, but accounting for ligands that act via another mechanism may require additional conformational states.

Going one step further than binary classification, we successfully quantify the prediction of E_max_ for the serotonin 3A receptor. A major hurdle in quantitatively predicting ligand efficacy has traditionally been the lack of large and consistently measured datasets. The available data is often scattered across multiple scientific publications, utilizing disparate assay conditions, different cellular contexts, and varying measurement methodologies. These inconsistencies make direct comparisons difficult and have often restricted predictions to qualitative, binary outcomes (agonist vs. antagonist). Using a consistent dataset on a system with a well characterized conformational spectrum,^43,66^ we demonstrate that binding free energy can serve as a quantitative predictor of a ligand’s agonism, given sufficient knowledge of a target’s activation mechanism. This finding further validates the underlying two-state model and demonstrates that our computational approach is suitable for the quantitative prediction and detailed mechanistic interrogation of ligand efficacy.

Membrane-facing binding pockets, such as those in TRPML1 and KCNQ2, present a particular challenge: The surrounding lipid environment must be carefully managed, as poorly equilibrated or incorrectly positioned lipids can introduce instabilities that compromise prediction reliability. One approach is to seed simulations with multiple independent lipid configurations, a simple version of which we used for TRPML1 when running once with the experimentally resolved lipid and once without. Although individual runs from this set may be less converged in isolation, the approach proved successful in aggregate. Beyond perfect separation of agonists and antagonists, this approach also achieved good agreement for the maximum response measured for the agonists. For KCNQ2, pre-equilibrating the membrane produced stable trajectories and consistent lipid positions. An important caveat: The ligands examined in the successful prediction on TRPML1 were structurally similar and occupied closely related poses, meaning that lipid configurations did not diverge substantially between compounds. For ligands from a new chemical series which may adopt distinct poses or displace lipids differently, greater caution is warranted in these membrane-facing sites than in the water-exposed or occluded binding pockets encountered in the extracellular domains. It is likely that these insights translate well to membrane-facing binding pockets of other membrane proteins like GPCRs or transporters, since they are conceptually similar across target families. We thus advise that accurate free-energy calculations for membrane-facing binding pockets in general require deliberate treatment of the lipid environment as a major consideration, not merely a boundary condition.

KCNQ2 represents a more challenging case in which agonists and antagonists separate along ΔΔG but the threshold shifts by 2 kcal/mol from the theoretical zero (Figure 4B/D). Since functional data was available only for a small number of ligands and from disparate sources, we have to be cautious when interpreting the results in search of potential causes. First, we cannot exclude experimental or modeling artifacts, similar to previous issues with an unresolved loop in the serotonin 2A receptor,^12^ and with the way electrostatic polarization effects are incorporated as zero-order bonds in simulations of integrin αIIbβ3.^14^ For this membrane-exposed binding pocket, the lipid environment is a potential factor, though we consider it unlikely to be responsible here due to the careful equilibration. A more fundamental consideration is that KCNQ2, as a voltage-gated channel, does not activate through conformational change alone.^76^ Besides effects of their structural preference, ligands could also modulate the likelihood of activation by a local change of the dielectric environment. The ligands in our study are all neutral though and we would expect such effects mostly from charged ligands like poly-unsaturated fatty acids.^77^ As an additional challenge for bridging structural modeling and pharmacology, KCNQ2 and KCNQ3 typically assemble as heteromers,^70^ and modeling based on the structure of KCNQ2 resolved in isolation may omit contributions from the KCNQ3 subunit to the binding pocket geometry and gating response. Despite these uncertainties, the successful classification using a shifted threshold demonstrates the practical utility of separating ligands based on their functional response, even in challenging systems that deviate from the ideal two-state model.

Despite the good performance demonstrated in our six examples, caution is always necessary as challenges may still arise even when formal requirements for functional response modeling are met, i.e., when functionally distinct conformational states and reliable ligand poses are available. One potential complication is that the two structures may not fully represent the structural landscape, potentially requiring more structures or looser restraints. The systematic approach of deriving restraints from unbiased molecular dynamics (MD) simulations, previously employed on GPCRs,^12^ was not used here because MD simulations of the entire ion channels are more costly. Another problem, here demonstrated on KCNQ2, is an imperfect threshold of the resulting separation of agonists and antagonists along ΔΔG. If the threshold is known, the classification can still be excellent. Since the model should ideally be tested on a few known ligands anyway, the threshold can be determined in this process. Finally, some ligands exhibit more complicated mechanisms. As an example from this work, CBD and HN37 each occupy two distinct but adjacent binding sites per subunit. While modeling the binding of only one molecule to the core site yielded good classification in this case, it was insufficient to capture the full details of each ligand’s respective action on the target. Even though it worked on these two ligands, it may not generalize to ligands with other pairing mechanisms or pairwise binding to other targets. Therefore, for real-world drug discovery, it is recommended to test the underlying structural hypothesis before scaling to large datasets, to be cautious when generalizing to new ligand types, and to regularly update the model with new insights.

Beyond its utility for known structures, AB-FEP offers a framework for validating the model quality of experimentally determined structures and of mechanistical hypotheses. Assume that we have a set of ligands with known functional response from a reliable assay but a set of structures fails to show the expected thermodynamic preference for the known ligands. This may indicate inaccuracies in one or more of the underlying structural models. It can even challenge the hypothesis about the underlying activation mechanism itself. We caveat that this approach is primarily effective for evaluating conformations within the binding pocket, which we discuss above as the main relevant region, and complementary methods are required to probe long-range effects. Still, such an objective assessment of practically applicable predictive power is a valuable check, especially for structures from electron cryo-microscopy, which lack the definitive mathematical solutions of X-ray crystal structures and whose quality and interpretability can remain subjective. The comparison of AB-FEP with functional assays then provides a thermodynamic benchmark to ground structural interpretations in functional reality.

In conclusion, we successfully apply FEP-based functional response modeling to a diverse range of ion channels, a challenging class of drug targets due to their complex architecture and multiple regulatory mechanisms. We demonstrate accurate binary classification across targets with both large and subtle conformational differences, utilizing appropriate position restraints, and achieved quantitative prediction of maximum response under near-ideal structural coverage and functional data availability. These successes reaffirm the fundamental thermodynamic principle that connects ligand efficacy to the binding free energy to different conformational receptor states. Furthermore, we address the specific challenge of membrane-facing binding sites, showing that with careful, deliberate treatment of the lipid environment we can achieve accurate prediction even for these challenging cases. The primary lesson learned is that FEP-based functional response modeling is a powerful and effective tool for high-accuracy prediction and mechanistic interrogation, particularly for selecting ligands with the correct function after initial binding screens. Ultimately, we establish a robust framework for integrating FEP calculations into ion channel drug discovery and propose target-specific adaptations for common scenarios.

## Methods

### General Strategy

As template structures for ligand efficacy prediction on ion channels, we selected at least two conformations, representing conducting/“active” and non-conducting/“inactive” states. Target selection prioritized a balance between broad coverage of diverse examples and data availability. For our simulations, we selected a stable sub-system of each target (Figure 1C), encompassing at least the ligand-binding domain(s). Each structure was prepared using the proteinprep workflow in Maestro,^78^ initially retaining the co-resolved ligand which was subsequently replaced for the FEP calculations.

For each target, we selected a set of ligands with known functional activity, giving preference to compounds for which co-crystallized structures or known binding poses were available. We sourced the functional annotations from the GtoPdb^35^ and from target-specific primary literature. The final binding poses were determined through three methods: extraction from solved experimental structures, modification of existing structures to model similar ligands, or docking using Glide. In the case of multiple reasonable candidate poses, we ran AB-FEP on each of them and used the lowest free energy value, i.e. the pose with the strongest predicted binding affinity. Similarly, we treat multiple reasonable lipid configurations (experimental vs. modeled lipids in TRPML1, six different modeled lipid configurations in KCNQ2) by picking the lowest free energy value.

For each combination of target, state, ligand, pose, and (where applicable) lipid configuration, we performed an AB-FEP calculation in FEP+,^4^ incorporating custom restraints. The restraints for ion channels are strong harmonic or flat-bottom harmonic position restraints with a force constant of 10 kcal mol^−1^ Å^−1^, centered on the experimental structure. The subset of restrained atoms and the width of the flat bottom (0 in case of harmonic restraints) vary by system. In previous work on GPCRs, restraint centers and widths for GPCRs were determined in a residue-specific way from unbiased molecular dynamics (MD) simulations.^12^ However, running MD on the entire large ion channel system would be computationally expensive, and simulating a smaller subsystem could introduce artifacts due to system instability. Furthermore, unlike nuclear receptors in previous work, the binding pocket architecture for most ion channel targets (except GluA2) is only subtly different between the agonist-bound and antagonist-bound states, meaning unrestrained simulations could lead to an undesirable overlap or convergence of the structural ensembles. The general strategy and settings described above were applied to each system unless explicitly noted otherwise in the following sections.

When possible, we tested the quantitative thermodynamic relationship between ligand efficacy and state-specific binding free energies, going beyond binary classification. In an ideal two-state system (“active”/“inactive”), the probability ratio between the two states is given by the Boltzmann factor:

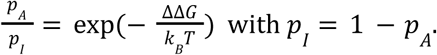

Solving for the probability of the active state yields a logistic function:

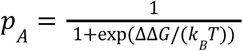

Assuming that the signal is proportional to the probability of the channel being in the active state, we fit to this relationship with an empirical offset *a* on ΔΔG as the sole fitting parameter:

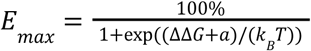

The fitting parameter accounts both for potential systematic shifts in the free energy calculations as well as for a potential threshold in the signal’s sensitivity. Properly testing this relationship requires experimental data for the maximum response from a consistently measured assay. Such data was available for two of the investigated receptors: 5-HT_3A_R and TRPML1.

### GluA2

For GluA2, we use the closed-state homo-tetramer in complex with the competitive antagonist ZK (fanapanel) from PDB 8SS6.^37^ For the agonist-bound state, we used the GluA2 ligand-binding domain in complex with glutamate from PDB 5NS9.^79^ From both structures, we only use the ligand-binding domain. Since the binding pocket of GluA2 differs strongly between the two states (Figure 2A), we employ flat-bottom harmonic restraints with a bottom width of 0.2 Å in one set of AB-FEP simulations and no restraints at all in another set. We probed all GluA2 agonists and antagonists available in the GtoPdb,^35^ as well as quisqualic acid as an additional agonist.^80^ Among the antagonists, the following were previously determined to be non-competitive: 4-BCCA,^81^ GYKI-53655,^82,83^ GYKI-53784,^83^ and talampanel.^84^ We include them to additionally probe whether our workflow can distinguish them correctly from the competitive antagonists that actually bind to the investigated binding pocket. See SI Table S1 for a full list of ligands. The AB-FEP runs of 5 ns length were preceded by an initial unbiased MD simulation of 1 ns. All heavy atoms of the backbone were restrained to the starting structure using flat-bottom harmonic restraints with a bottom half-width of 0.2 Å.

### GABA_A_R ρ1

A variety of conformational states of the GABA_A_R ρ1 are available, from which we choose two template structures bound to the native agonist GABA and two structures bound to inhibitors, mainly differentiated by the position of Loop C (Figure 3A). One of the GABA-bound structures is in the primed state (PDB 8RH7),^40^ the other one in the desensitized state (PDB 8OP9).^38^ While they show no significant difference in Loop C, the position of Loop B in the desensitized state is roughly halfway in between the primed state and the inhibitor-bound states. One of the inhibitor-bound structures (PDB 8OQ7) was resolved with TPMPA,^38^ the other one (PDB 9FRB) with CGP36742.^39^ They differ in the orientation of Met177 which points toward the ligand in the TPMPA-bound structure and away from it in the CGP36742-bound structure. For all modeling, we only consider the extracellular domain of two monomers of the pentameric channel (chains A and B in the respective PDB structures) and the ligand binding pocket at their interface. For six GABA_A_R ρ1 ligands we use experimental poses: GABA (agonist, PDB 8OP9),^38^ (S)-GABOB (agonist, PDB 9FRH),^39^ (R)-GABOB (agonist, PDB 9FRI),^39^ THIP (antagonist, PDB 9FRE),^39^ TPMPA (antagonist, PDB 8OQ7),^38^ and CGP36742 (antagonist, PDB 9FRB).^39^ For the ten additional ligands with functional data in the GtoPdb (five agonists and five antagonists), we generated poses via docking using Glide. See SI Table S2 for a full list of ligands. For most ligands, we use only the top pose in each template structure, with one exception: For aza-THIP, we use the top three poses because the top poses in both receptor states were flipped compared to THIP. The AB-FEP runs of 5 ns length were preceded by an initial unbiased MD simulation of 1 ns. All heavy atoms of the backbone were restrained to the starting structure using harmonic restraints.

### ɑ3β4 nAChR

We started from a pair of extremely similar structures of α3β4 nAChR, one bound to the agonist nicotine (PDB 6PV7) and one bound to the antagonist AT-1001 (PDB 6PV8), resolved within the same study (Figure 3B) and both suggested to be in a desensitized state.^41^ Just like for GABA_A_ R ρ1, we only model the extracellular domains of two monomers (chains A and B in the respective PDB structures) and the ligand binding pocket at their interface. We assembled a dataset with ligands of known functional activity and one decoy channel blocker (18-MC), see SI Table S3 for a full list. Poses were determined via docking using Glide. Up to four top poses were manually selected depending on their conformational variability and docking scores. To initially determine whether the differences between the two states are significant and functionally relevant and whether our standard simulations of 5 ns are sufficient to distinguish the only slightly different agonist-bound and antagonist-bound states, we performed an uncertainty analysis with four repeat simulations using different random seeds on a subset of five selected ligands: the antagonists AT-1001 and SR-16584 as well as the agonists nicotine, cytisine, and epibatidine. Based on the results (SI Figure S4), we decided to run the FEP stage of our simulations on the full dataset with 10 ns as additional precaution while keeping the initial unbiased MD stage at 1 ns. All heavy atoms of the backbone were restrained to the starting structure using harmonic restraints. Because of the very similar setup of both simulations, we used α3β4 nAChR to investigate the relation of the uncertainties of individual FEP calculations to the uncertainty of the difference ΔΔG as described in the Supplementary Text.

### 5-HT_3A_R

We picked two structures of the 5-HT_3A_R with extreme positions of Loop C, whose detailed conformations during inhibition and activation have recently been resolved structurally.^42,43^ Specifically, we used the non-conducting state from PDB 6W1Y, resolved with palonosetron,^42^ and the open-like receptor state from PDB 8FSB, resolved with serotonin (Figure 3C).^43^ In line with the other two Cys-loop receptors, we only consider the extracellular domains of two monomers (chains A and B in the respective PDB structures) and the ligand binding pocket at their interface. We probed all ligands from a consistently determined E_max_ dataset,^43,66^ see SI Table S4 for a full list. For all of them, we used the poses determined via Cryo-EM,^43^ or, if not directly determined, aligned the ligands to a resolved congeneric ligand. Since we expected the quantitative resolution of functional response aimed for in this study to be less error-tolerant, we extended the initial unbiased MD to 2 ns and the AB-FEP stage to 20 ns. All heavy atoms of the backbone were restrained to the starting structure using flat-bottom harmonic restraints with a bottom half-width of 0.25 Å.

### TRPML1

From a systematic study of TRPML1 activation,^44^ we chose the structure resolved with antagonist compound 9a (PDB 9HLA) and the one resolved with agonist compound 5 (PDB 9HJ8), both of which were co-resolved with a lipid, 3-sn-phosphatidylcholine, adjacent to the ligand binding pocket (Figure 4A). For this channel, we include the transmembrane domain of all four subunits in the simulated sub-system with the exception of the transmembrane helices S1 and S2. AB-FEP was performed once with the co-resolved lipid removed and once keeping it in place. In the latter case, we slightly adapted the lipid tails to accommodate all investigated ligands within a single AB-FEP run. We probed all ligands included in the same structural study (full list in SI Table S5) and used either their resolved poses or, if not resolved, aligned them to a similar ligand. To allow more time to sample lipid rearrangements, we extend the standard length of the FEP stage from 5 ns to 10 ns, keeping the initial unbiased MD stage at 1 ns. Backbone atoms C, N and Cɑ were restrained to the starting structure using harmonic restraints.

### KCNQ2

From a study of the mechanism of a KCNQ2 antagonist,^45^ we chose the agonist-bound state resolved with Ebio2 (PDB 9IXY) and the antagonist-bound state resolved with Ebio3 (PDB 9IXZ), hoping to leverage the small difference in the binding pocket observed between those two states (Figure 4B). To not destabilize the lipid environment, we include the entire transmembrane domain in the simulations. To obtain an equilibrated membrane, we performed six independent MD runs from each state, with the protein backbone restrained. Lacking a large consistently measured dataset, we assembled ligands with known poses in the pore domain and with functional data from across multiple publications. Besides the two template ligands, these are: retigabine (RTG, agonist, PDB 7CR2),^30^ RTG-S1 (aligned to RTG), Ebio1 (PDB 8IJK), and Ebio1-S1 (PDB 8X43),^32^ pynegabine (HN37, agonist, PDB 8W4U) and cannabidiol (CBD, agonist, PDB 8J03),^31^ as well as QO-83 (agonist, PDB 8X01).^33^ As for TRPML1, we ran the FEP stage for 10 ns, keeping the initial unbiased MD stage at 1 ns. Backbone Cɑ atoms were restrained to the starting structure using flat-bottom harmonic restraints with a half-bottom width of σ = 0.5 Å for the agonist-bound state and σ = 1 Å for the antagonist-bound state.

The two ligands CBD and HN37 can bind both alone and as dimers.^31,32^ We ran additional AB-FEP simulations for a second molecule with the first one bound. In both cases, the second molecule clashed with lipids and HN37 additionally with the cofactor PIP_2_ in the open state (SI Figure S11A/B). We removed the clashing lipids and clashing PIP_2_ in those simulations, letting surrounding lipids relax in their place during an extended initial MD stage of 5 ns. As for the first molecules, we ran the FEP stage for 10 ns. Backbone Cɑ atoms were restrained to the starting structure using flat-bottom harmonic restraints with a half-bottom width of σ = 1.0 Å.

## Data Availability

Input poses and numerical AB-FEP results are available on GitHub: https://github.com/schrodinger/ion-channels-efficacy

## Acknowledgments

We thank Emelie Flood, Kevin DeMarco, and Jonas Kaindl for helpful discussions.

