## Supplementary Information for "State-specific binding thermodynamics predicts ligand efficacy across ion-channel families"

### Supporting Text

#### Probabilities as Classification Scores

Instead of using the free energy difference directly as a classification score, it can be instructive to also calculate the resulting probability $p_i$ for a ligand to be an agonist. The probability to be an antagonist is then simply $1-p_i$ This probability corresponds to the probability for the unknown true value $\theta_i$ of ΔΔG to be below the classification threshold τ, given the observed ΔΔG value $d_i$ and an assumed Gaussian uncertainty $\sigma_i$. We can obtain it by integrating the corresponding Gaussian distribution $N(d_i,\sigma_i^2)$ over the negative real numbers. This is equivalent to

$$p_i = \Phi\left(\frac{\tau - d_i}{\sigma_i}\right)$$

with Φ the standard-normal cumulative distribution function. These scores can be used to assess the confidence of the binary prediction. We can, for example, compare the individual scores to to the corresponding ground-truth sign $y_i$ which is –1 for agonists and +1 for antagonists by calculating the binary RMSE:

$$\varepsilon = \sqrt{\frac{1}{n}\sum_{i=0}^{n}(p_i - y_i)^2}$$

as a measure for the quality of the prediction on a dataset with $n$ ligands. We can also optimize the classification threshold τ by minimizing ε.

#### Probability of Correct Prediction

We now estimate the probability $q_i$ for our predictor to correctly classify ligand $i$. For simplicity, we further assume a classification threshold τ=0 as predicted by the simple two-state model, even though it can deviate in practice. For the true value $\theta_i$ of ΔΔG, our model produces the observed value $d_i$ according to the Gaussian distribution $N(\theta_i, \sigma_i^2)$. Note that, since we assume only stochastic sampling noise, the value $\theta_i$ may be different from the actual free energy difference. From this, we calculate the probability of recovering the correct sign analogous to $p_i$ above but include the ground-truth label $y_i$ to flip the sign accordingly. This yields

$$P(y_i d_i > 0 \mid \theta_i) = \Phi\left(\frac{y_i \theta_i}{\sigma_i}\right).$$

with Φ the standard-normal cumulative distribution. Since we don't know the true value of ΔΔG, our best guess is the observed value $d_i$ and its uncertainty $\sigma_i$. We can calculate the marginal probability as the expected value of the conditional probability calculated above:

$$q_i = E_{\theta_i | d_i, y_i}\left[\Phi\left(\frac{y_i \theta_i}{\sigma_i}\right)\right] = \int_{-\infty}^{\infty} \Phi\left(\frac{y_i \theta_i}{\sigma_i}\right) p(\theta | d_i, y_i)\, d\theta$$

The probability distribution of $\theta$ is the Gaussian likelihood we assume from the simulation, modulated by our prior expectation π.

$$p(\theta | d_i, y_i) \sim N(\theta_i, \sigma_i^2)\, \pi(\theta | y_i)$$

With the appropriate normalization, this leads to the following general expression

$$q_i = \frac{\int_{-\infty}^{\infty} \Phi\left(\frac{y_i \theta_i}{\sigma_i}\right) N(\theta_i, \sigma_i^2)\, \pi(\theta | y_i)\, d\theta}{\int_{-\infty}^{\infty} N(\theta_i, \sigma_i^2)\, \pi(\theta | y_i)\, d\theta}$$

If we don't know whether our simulations will always predict the correct labels except for the assumed stochastic noise, we have to use a flat prior

$$\pi(\theta \mid y_i) \sim 1$$

and if we assume that all $\theta_i$ values possess the correct sign, i.e. our simulations can be expected to produce the true experimental label on average, a Heaviside step function can be applied to restrict the prior to the appropriate half of the real numbers:

$$\pi(\theta \mid y_i) \sim H(y_i \theta).$$

Using a flat prior we get

$$q_i = \Phi\left(\frac{y_i d_i}{\sigma_i \sqrt{2}}\right)$$

where Φ is the standard-normal cumulative distribution function. And using the step prior, we get

$$q_i = \frac{1}{2} + \frac{\Phi(a_i/\sqrt{2})^2}{2\Phi(a_i)} \text{ with } a_i = \frac{y_i d_i}{\sigma_i}$$

These scores can be used to assess the confidence of the binary prediction. We can, for example, calculate the probability of a prediction with perfect accuracy $a$ on a dataset of n ligands as all correct

$$P(a = 100\%) = \prod_{i=0}^{n} q_i$$

### Propagation of Uncertainty to ΔΔG

The uncertainty σ of ΔΔG is derived from the uncertainties $\sigma_A$ and $\sigma_I$ of the individual values $\Delta G_A$ and $\Delta G_I$. If we assume the individual uncertainties to be uncorrelated, we calculate the total uncertainty according to

$$\sigma^2 = \sigma_A^{\ 2} + \sigma_B^{\ 2}$$

If we assume the total uncertainty to be a combination of statistic sampling uncertainty and systematic uncertainties from the simulation setup, we calculate it as

$$\sigma^2 = \sigma_A^{\ 2} + \sigma_B^{\ 2} - 2\rho\, \sigma_A \sigma_B + \sigma_{A,s}^{\ 2} + \sigma_{B,s}^{\ 2}$$

with the correlation factor ρ. Analysis of the results for α3β4 nAChR (Figure S5) suggests that the systematic uncertainties from the system setup largely cancel each other out when the differences between the states are small. In some cases, they do not cancel each other out completely but the remaining artifact is the same for all (or a subset of) ligands. Then it contributes to a shift of the classification threshold away from the theoretical zero. Larger differences between states may make it necessary to also adapt protonation states or force field parameters which can introduce systematic deviations. However, stronger differences between the states can make classification easier overall (if we adapt the threshold accordingly).

### Supporting Figures

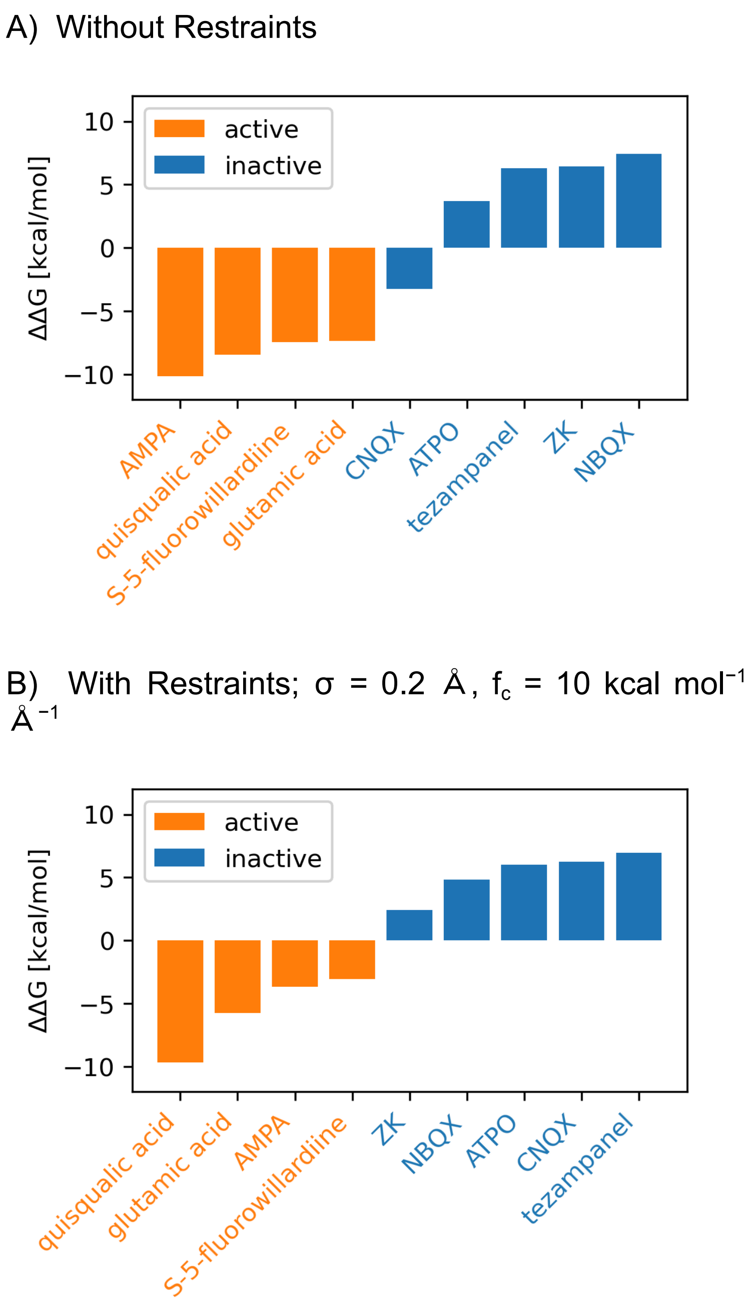


**Figure S1**: Comparison of functional predictions via $\Delta\Delta G = \Delta G_A - \Delta G_I$ for GluA2 binders with and without restraints. The bars and names of active compounds are shown in orange and those of inactive compounds in blue. Failed simulations and positive predicted values of $\Delta G_{I/A}$ are represented as 0 kcal mol$^{-1}$. Numerical results are listed in Table S1.

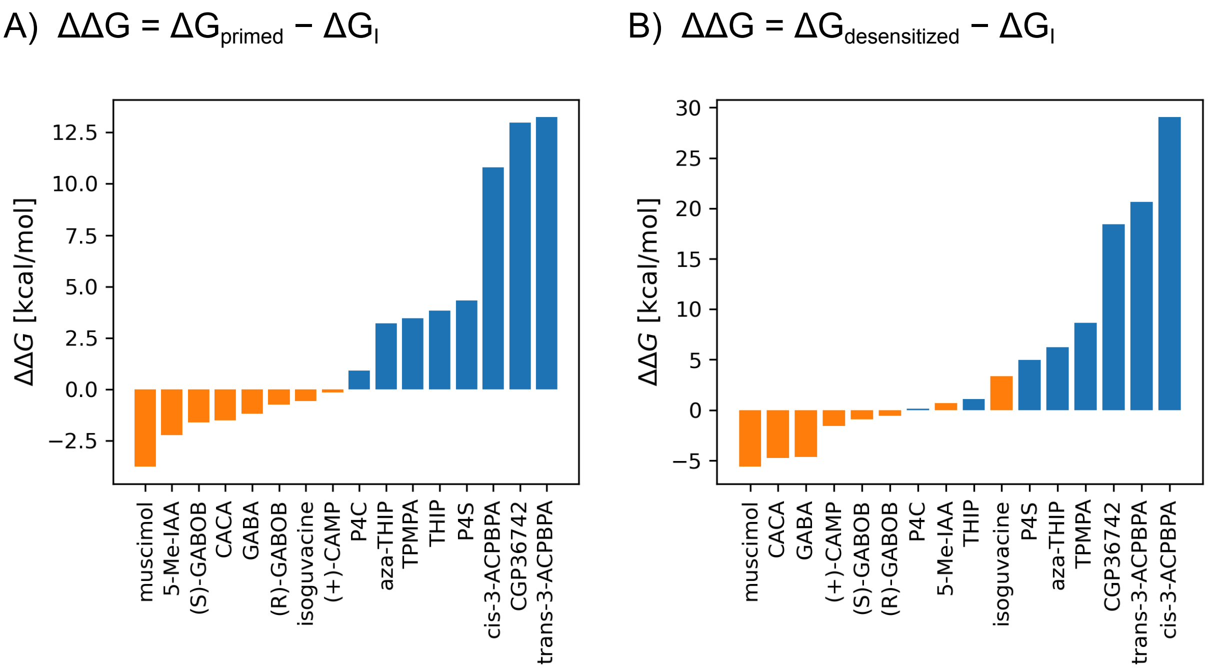


**Figure S2**: The desensitized conformation alone is a slightly worse representation for the active state than the primed conformation alone. (A) Prediction results using $\Delta G_{primed}$ as $\Delta G_A$, with perfect separation of agonists and antagonists. (B) Prediction results using $\Delta G_{desensitized}$ as $\Delta G_A$, with two wrong predictions and generally weaker separation.

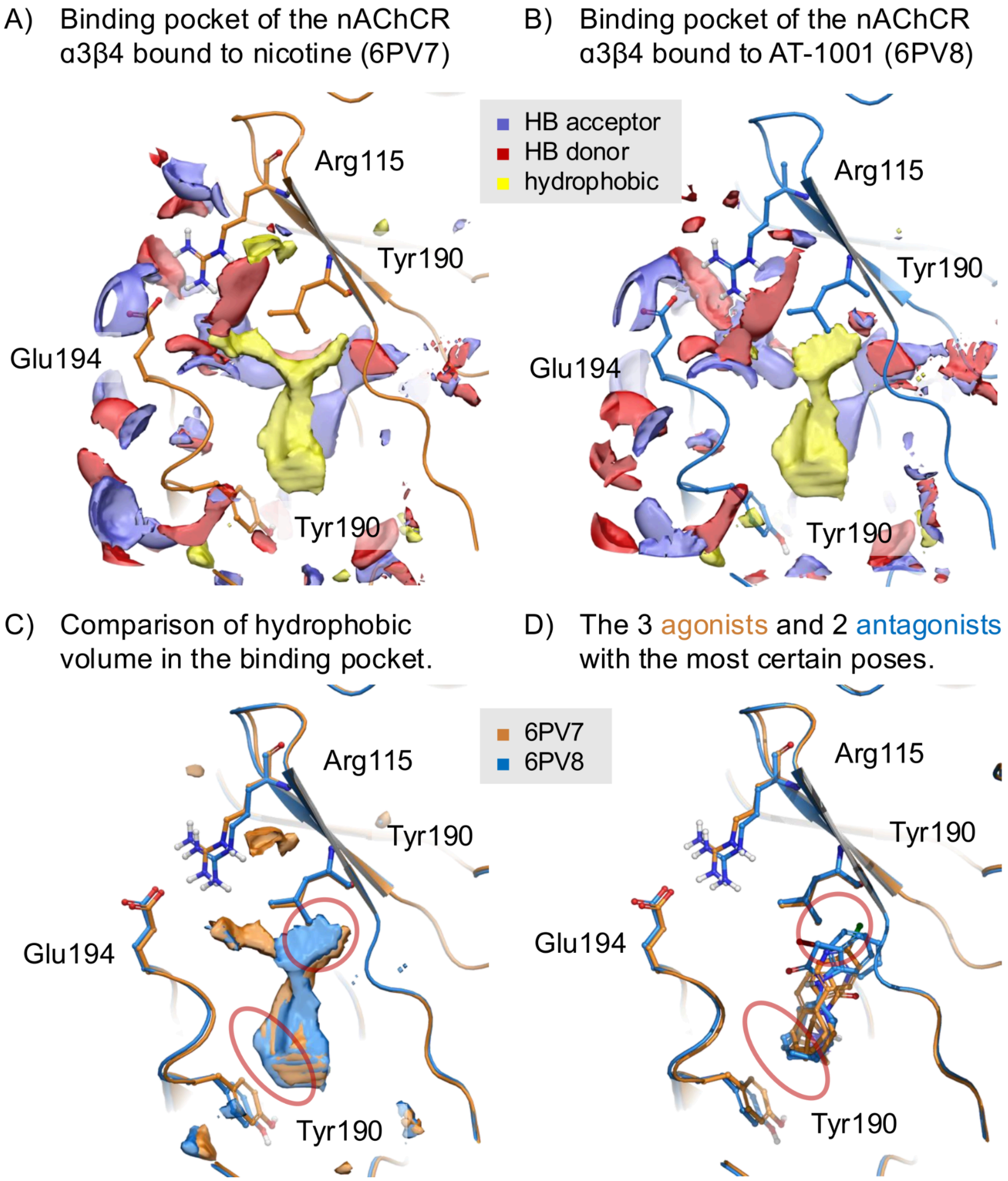


**Figure S3**: Restraining small differences in the α3β4 nAChR has outsized influence on the accessible volume in the binding pocket of the AB-FEP simulations. We show the volume of the binding pocket as calculated via Schrodinger SiteMap for (A) the α3β4 nAChR in the agonist-bound state (PDB 6PV7, orange) and (B) in the antagonist-bound state (PDB 6PV8, blue). The backbone structures are shown as ribbons. (C) The hydrophobic volume in both states (6PV7 orange, 6PV8 blue) with the most striking differences highlighted using red ovals. (D) The agonists nicotine, cytisine and epibatidine and the antagonists AT-1001 and SR-16584 with the regions corresponding to the most important volume differences highlighted in red circles. Other ligands showed multiple reasonable candidate poses in docking.

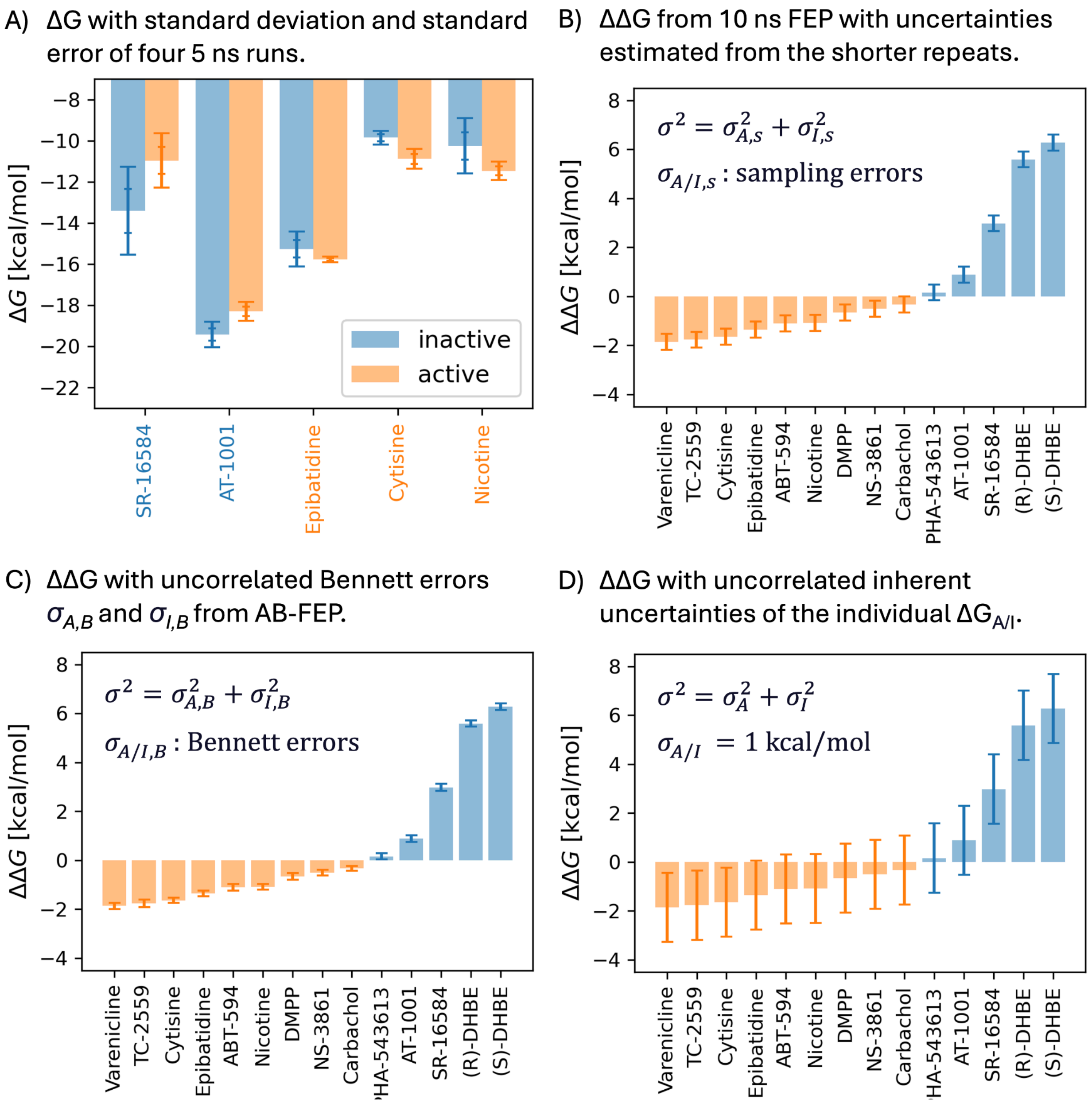


**Figure S4**: (A) Predictions using the average ΔG of four 5 ns repeat runs. The binding ΔG to the inactive state of antagonists AT-1001 and SR-16584 is correctly predicted to be stronger, and weaker for the three agonists. Standard errors do not overlap, confirming a significant link between the small structural differences in the binding pocket and the ligand function. Some standard deviations overlap, which means that a single 5 ns run per ligand may not always yield correct predictions. (B) ΔΔG for all α3β4 nAChR ligands from single 10 ns AB-FEP runs with uniform uncertainties estimated from the standard deviation of the shorter repeats as the median standard deviation scaled by √2. (C) ΔΔG for α3β4 nAChR ligands with the uncorrelated Bennett errors σA,B and σI,B from AB-FEP as the only uncertainties. (D) ΔΔG for α3β4 nAChR ligands with the uncorrelated inherent uncertainties of the individual ΔGA/I values from AB-FEP.

A) Probabilities of no or at most one wrong classification for the α3β4 nAChR ligands as a function of the error correlation coefficient, assuming Bennett errors as the only source of statistical uncertainty.

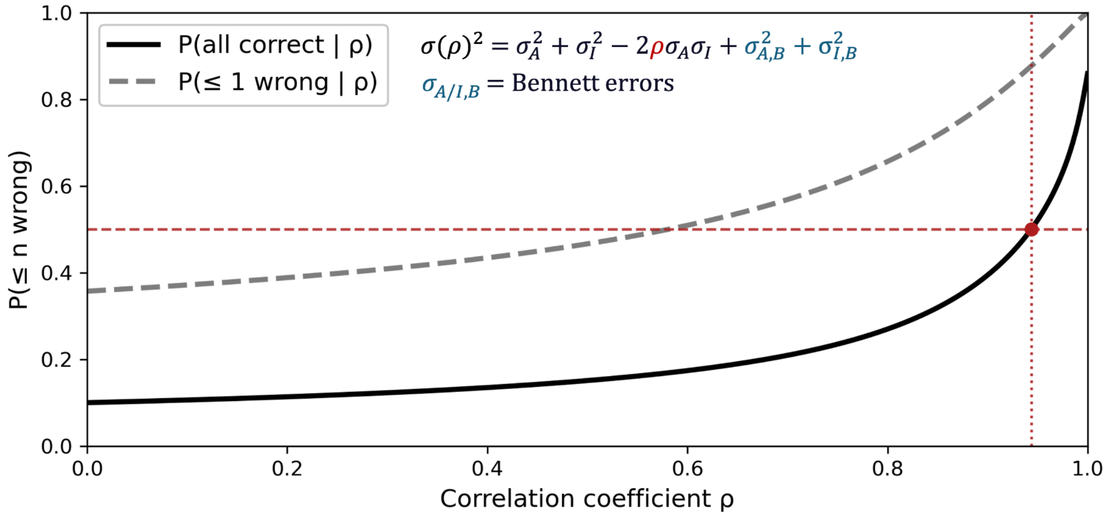


B) Probabilities of no or at most one wrong classification for the α3β4 nAChR ligands as a function of the error correlation coefficient, using a uniform uncertainty estimated from shorter repeat runs.

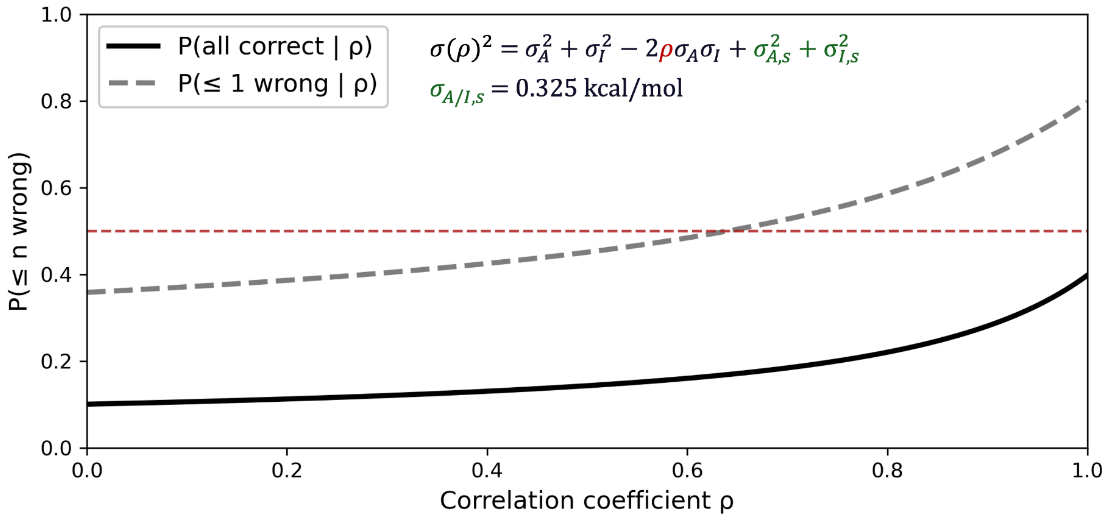


**Figure S5**: Uncertainty correlation for α3β4 nAChR. Probabilities of three different levels of prediction performance over the correlation coefficient between the individual uncertainties, calculated as derived in the Supplementary Text for (A) Bennett errors estimated from the simulation trajectory alone as the source of statistical noise and (B) uncertainty estimates from repeat runs of 5 ns as in Figure S4B.

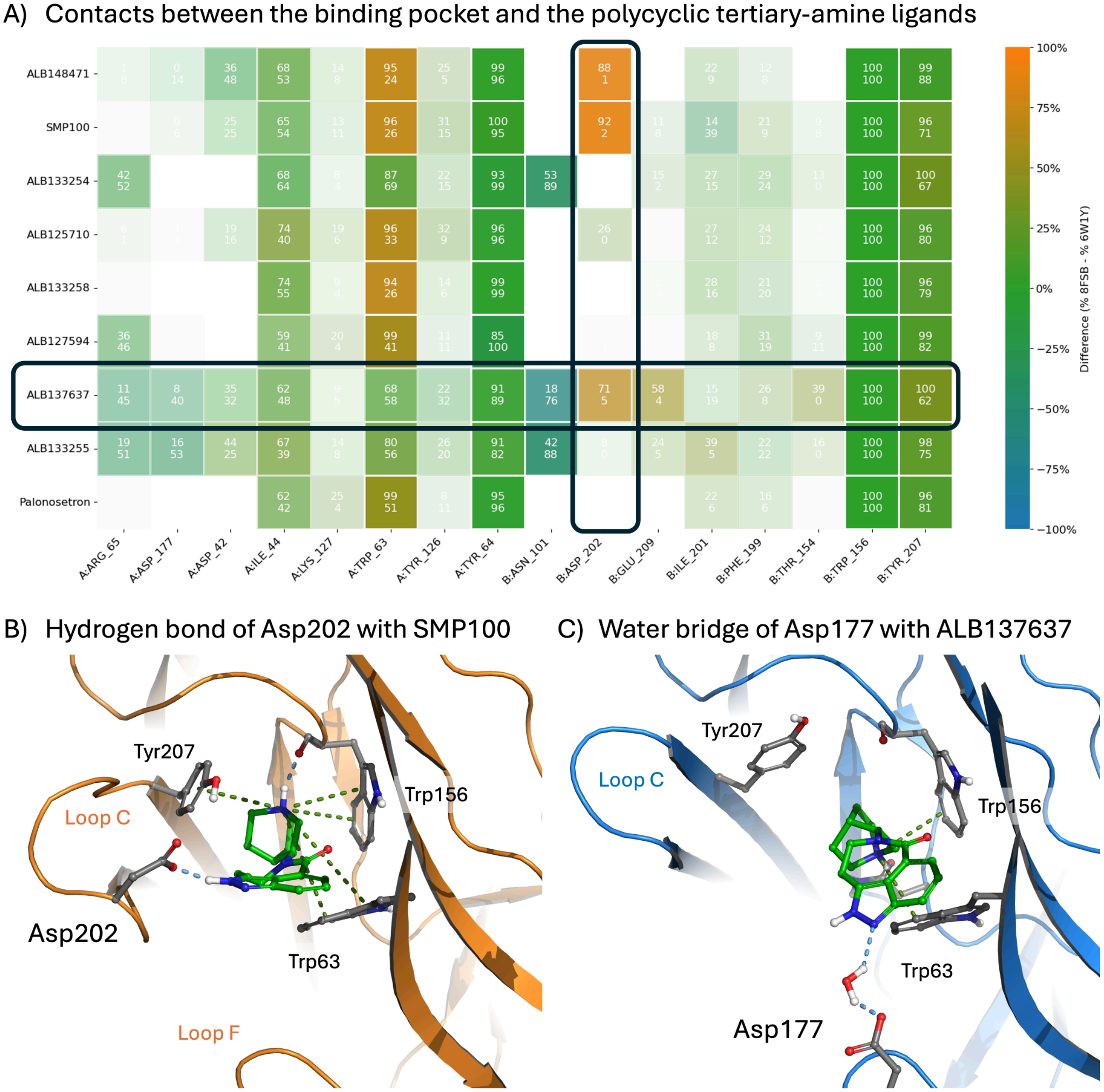


**Figure S6**: Interaction Comparison for 5-$HT_{3A}$R (A) Comparison of interactions between the binding pocket of 5-$HT_{3A}$R with the polycyclic tertiary-amine ligands that form a congeneric series. (B) Representative FEP frame of SMP100 in the binding pocket of 5-$HT_{3A}$R from PDB 8FSB. The hydrogen bond with Asp202 can only be formed by agonists, with the exception of ALB137637 which is an enantiomer of SMP100. (C) Representative FEP frame of ALB137637 in the binding pocket of 5-$HT_{3A}$R from PDB 6W1Y. Due to the different conformation of stereocenters (SRS instead of RSR), the charged amine anchors ALB137637 differently in the pocket, allowing a water bridge to Asp177. The free energy reduced by the hydrogen bond to Asp202 in the open-like state is at least partially offset by the formation of this water bridge. This offers a possible explanation for this ligand's reduced agonism despite its ability to form the hydrogen bond with Asp202.

A) Comparison of ΔG from binding experiments and AB-FEP.

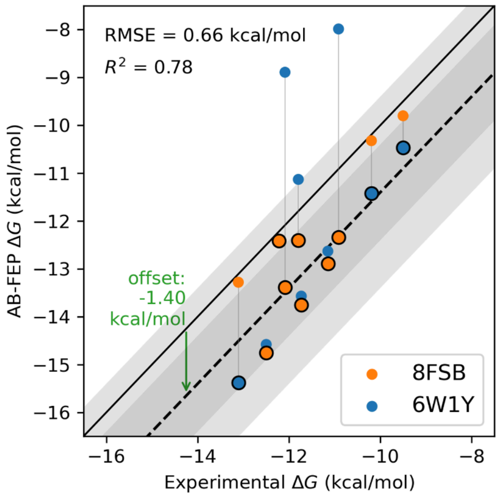


B) Residues close to the binding pocket that differ between mouse and human $5HT_{3A}R$

| Chain | 8FSB residue | Loop | Human ortholog residue | Distance to serotonin | Change type |
|---|---|---|---|---|---|
| A | Ile201 | C | Met223 | 3.8 Å | conservative, hydrophobic |
| A | Asp202 | C | Glu224 | 5.5 Å | conservative, acidic |
| A | Ile203 | C | Ser225 | 9.4 Å | non-conservative, hydrophobic → polar |
| A | Ser206 | C | Tyr228 | 7.5 Å | non-conservative small polar → bulky aromatic |
| E | Met45 | D | Val67 | 8.7 Å | conservative, hydrophobic |
| E | Ile180 | F | Val202 | 6.8 Å | conservative, hydrophobic |

C) $5HT_{3A}R$ in PDB 8FSB (mouse).

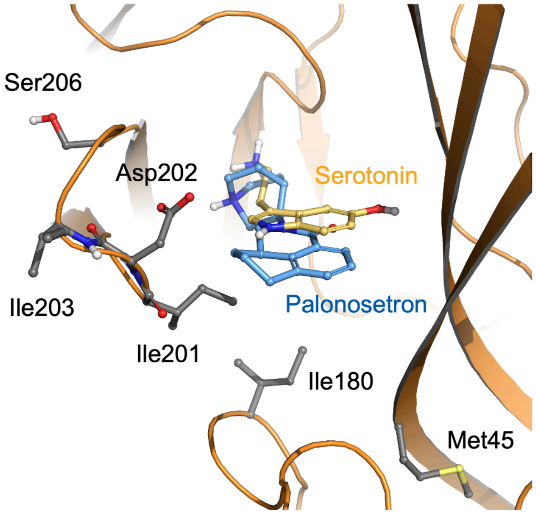


D) $5HT_{3A}R$ in humanized PDB 8FSB.

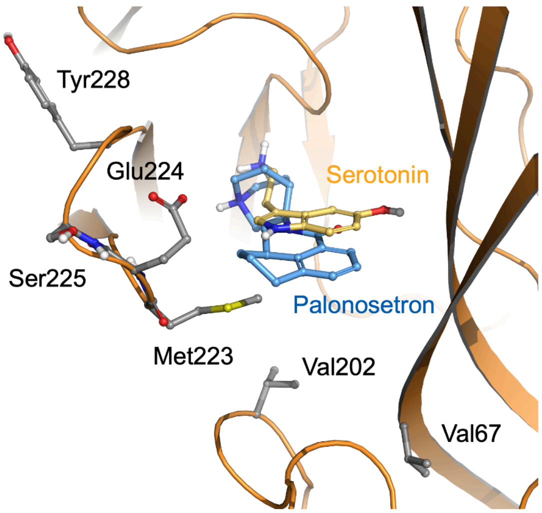


**Figure S7**: (A) Correlation between computational and experimental binding free energy of 5-$HT_{3A}R$ ligands. The lowest computational value of ΔG for each ligand (circled in black) was used, leading to a better correlation than each structure alone. The offset was determined via a numerical fit. (B) Residues close to the binding pocket, defined as within 10 Å of serotonin in 8FSB, that differ between mouse and human 5-$HT_{3A}R$. (C) Residues in original 8FSB that differ between mouse and human 5-$HT_{3A}R$. Palonosetron from aligned 6W1Y. (D) Residues in humanized 8FSB that differ between mouse and human 5-$HT_{3A}R$. Palonosetron from aligned 6W1Y.

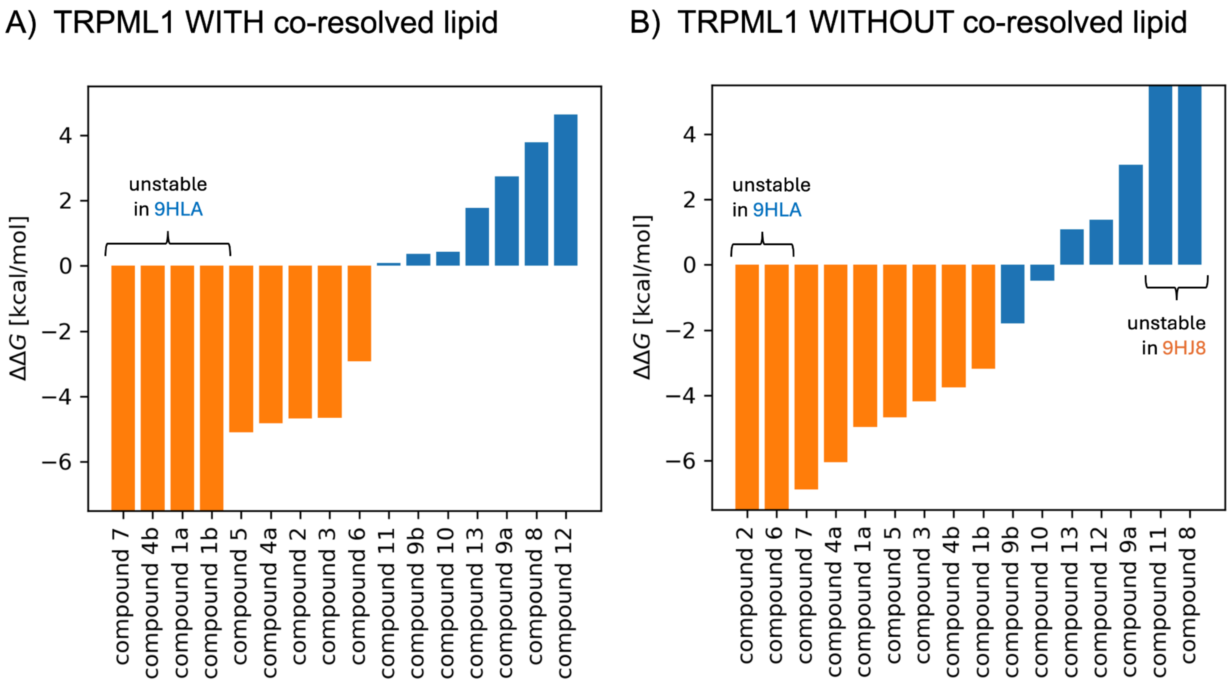


**Figure S8**: Functional response predictions for TRPML1 ligands from AB-FEP simulations with experimentally resolved lipids (left) and without (right). ΔΔG is the difference of the binding free energy $\Delta G_A$ calculated on the active state of TRPML from PDB 9HJ8, and the binding free energy $\Delta G_I$ calculated on its inactive state from PDB 9HLA. Unstable simulations are counted as ΔG = 0, resulting in a prediction strongly favoring the other state.

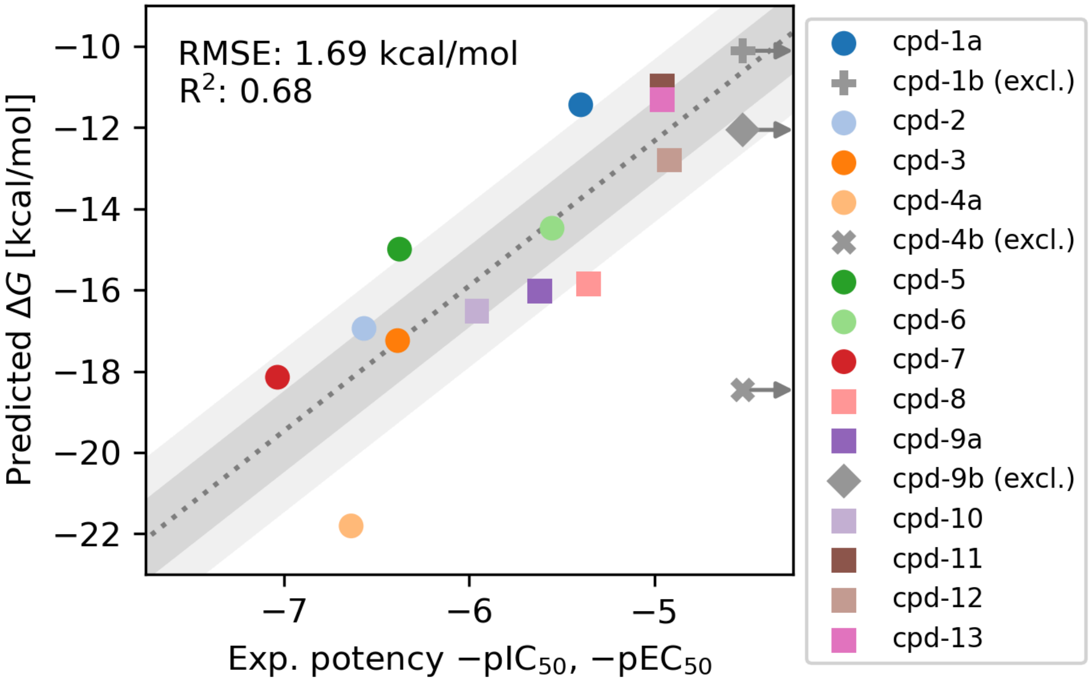


**Figure S9**: Correlation between the predicted binding free energy to TRPML1 with the experimentally determined functional potency −pIC50 for antagonists and −pEC50 for agonists, an approximate but imperfect proxy for the binding affinity where −pX50 = log10(X50/M). Agonist EC50 values were obtained by applying each test compound alone and normalizing channel activation to the maximal response produced by 10 µM ML-SA5 while antagonist IC50 values were instead measured as inhibition of channel current pre-activated with an EC70 concentration of ML-SA5 and normalized between the ML-SA5 reference current and complete block by lanthanum [Reeks et al., Structure 33, 1374–1385, 2025]. Thus, EC50 and IC50 describe different functional experiments and should not be interpreted as directly equivalent binding constants. Furthermore, the ligand-binding pocket faces the membrane and ligand partitioning into the membrane was not included in AB-FEP, so systematic or ligand-dependent offsets in the absolute affinity values are expected. This comparison should therefore be interpreted as an assessment of the qualitative trend rather than a quantitative validation of absolute binding affinity. Low-potency compounds (EC50, IC50 > 30 µM) are shown in gray at the upper bound of their pIC50 or pEC50 values and excluded from the fits.

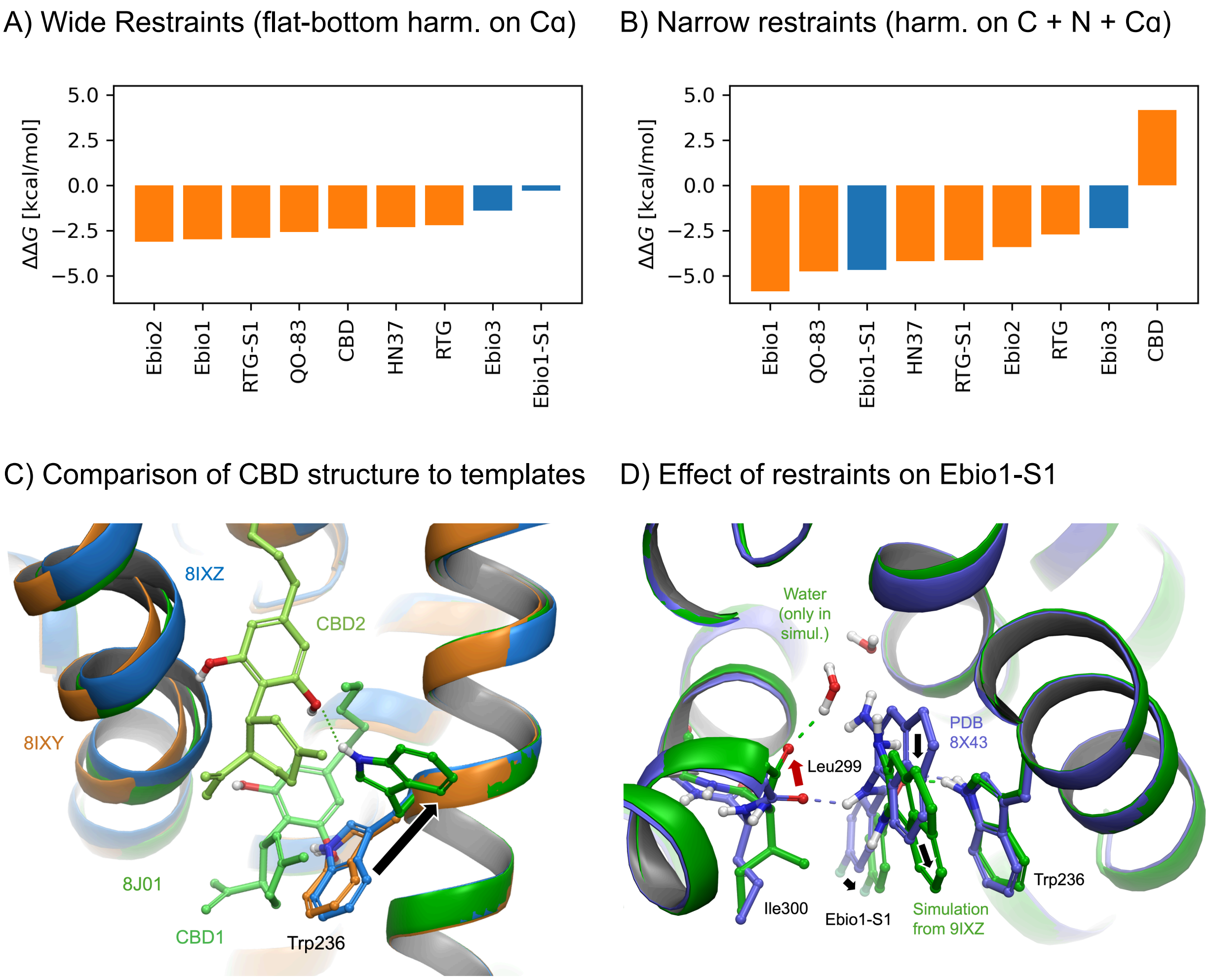


**Figure S10**: Narrower position restraints in AB-FEP simulations of KCNQ2 decrease the classification performance. **(A)** Results with wide restrains, as in Figure 4D. **(B)** Results with narrower restraints: harmonic restraints on backbone atoms C, N, and Cα. All antagonists are predicted as agonists and the only ligand predicted to be an antagonist is actually an agonist (CBD). A few relative trends within groups of congeneric ligands are represented correctly: ΔΔG is higher for antagonist Ebio3 than for agonist Ebio2.[44] Ebio1-S1 shifts up ΔΔG compared to the channel-opening Ebio1 from which it was derived.[31] Similarly, RTG has a higher ΔΔG than its more opening derivative RTG-S1 where a methyl substitution of an amino group disrupts the hydrogen bond interaction with S303 and F305 of KCNQ2,[31] and a higher ΔΔG than QO-83 which is more opening due to its additional cyclopentyl group forming hydrophobic interactions with V225.[32] **(C)** Comparison of an experimental structure of KCNQ2 bound to CBD (8J01, green) to the active-state (8IXY, orange) and inactive-state (8IXZ, blue) structures. While the backbone of the CBD-bound structure is similar to the active state, CBD binds in pairs and Trp236 is flipped compared to both other structures. The CBD structure is outside the range represented in the strongly restrained AB-FEP simulations. **(D)** Ebio1-S1 needs subtle backbone rearrangements that are prevented by the narrow restraints. Comparison of a representative structure from an inactive-state AB-FEP simulation with narrow restraints (green) to the experimental structure of KCNQ2 bound to

Ebio1-S1 (8X43, purple). To accommodate Ebio1-S1, the backbone oxygen of Leu299 needs to shift inward (red arrow) together with neighboring backbone atoms. In the simulation, these changes cannot happen because of the narrow restraints, resulting in the ligand to shift outward (black arrows) compared to its experimental pose into a less favorable pose while its hydrogen bond to Leu299 is taken over by a water molecule.

A) Template clashes with second CBD

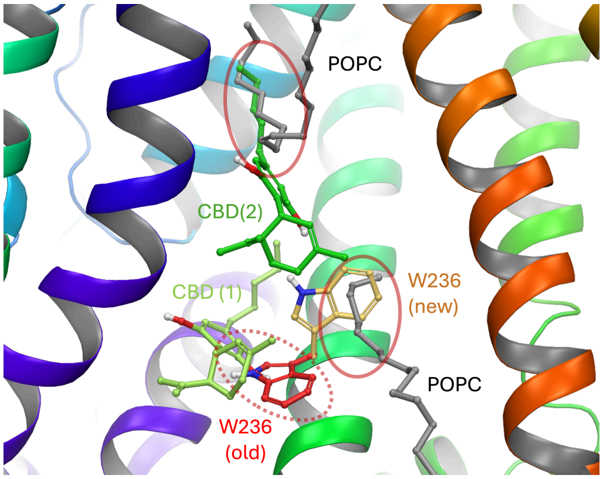


B) Template clashes with second HN37

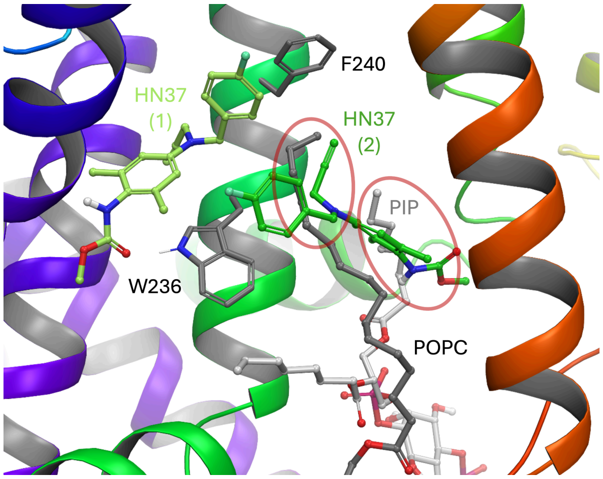


C) Binding results of second CBD and HN37 do not fundamentally change the total ΔΔG.

| Ligand | CBD | HN37 |
|---|---|---|
| $\Delta G_{A,2}$ [kcal/mol] | −17.0 | −14.7 |
| $\Delta G_{I,2}$ [kcal/mol] | −17.5 | −14.6 |
| **$\Delta\Delta G_2$ [kcal/mol]** | **0.5** | **−0.1** |
| ΔΔG [kcal/mol] | −2.4 | −2.3 |
| $\Delta\Delta G + \Delta\Delta G_2$ [kcal/mol] | −1.9 | −2.4 |

D) Functional states of resolved structures illustrate different mechanisms.

| Ligands in Sample | Ligands in Structure | State of the Gate |
|---|---|---|
| CBD | 2 CBDs | closed |
| **CBD, $PIP_2$** | **1 $PIP_2$, 2 CBDs** | **open** |
| CBD, $PIP_2$ | 1 $PIP_2$, 1 CBDs | open |
| HN37 | 2 HN37s | closed |
| **HN37, $PIP_2$** | **2 HN37s** | **closed** |
| HN37, $PIP_2$ | 1 $PIP_2$, 1 HN37 | open |

Select structures from Ma et al., Nat. Commun. (2023)14:6632

**Figure S11**: The role of second molecules of CBD and HN37 in KCNQ2. (A) The second copy of CBD binds within the same groove as the first one and clashes with one lipid in the template setup and the rotation of W236 observed in AB-FEP of the first copy clashes with another one (red circles). We remove both lipids in the new simulation. (B) The second copy of HN37 displaces $PIP_2$ and one POPC (red circles), both of which we remove in the new simulation. (C) Binding affinities of both ligands differ only slightly between the two states (CBD by 0.5 kcal/mol, HN37 by −0.1 kcal/mol), not fundamentally changing the prediction result. (D) Structures resolved with each ligand alone as well as with the cofactor $PIP_2$ show that the second HN37 molecule acts antagonistically not by shifting the conformational equilibrium but by displacing the necessary cofactor $PIP_2$. This effect is not reflected in our workflow.

### Supporting Tables

| Ligand | $\Delta G_A$ unrestr. | $\Delta G_I$ unrestr. | $\Delta G_A$ restr. | $\Delta G_I$ restr. | Min. $\Delta G_I$ | $\Delta\Delta G$ restr. | Activity |
|---|---|---|---|---|---|---|---|
| S-5-fluorowillardiine | −14.5 | −7.1 | −10.8 | −7.8 | −7.8 | −3.0 | agonist |
| NBQX | NaN | −7.4 | NaN | −4.8 | −7.4 | 4.8 | antagonist |
| tezampanel | NaN | −6.3 | NaN | −6.9 | −6.9 | 6.9 | antagonist |
| ZK | −0.4 | −6.9 | >0.0 | −2.4 | −6.9 | 2.4 | antagonist |
| CNQX | −9.9 | −6.7 | >0.0 | −6.3 | −6.7 | 6.3 | antagonist |
| quisqualic acid | −14.5 | −6.1 | −16.2 | −6.6 | −6.6 | −9.7 | agonist |
| ATPO | NaN | −3.7 | NaN | −6.0 | −6.0 | 6.0 | antagonist |
| AMPA | −16.5 | −6.5 | −9.8 | −6.1 | −6.3 | −3.7 | agonist |
| glutamic acid | −12.1 | −4.7 | −11.2 | −5.4 | −5.4 | −5.7 | agonist |
| 4-BCCA | −2.7 | −2.2 | >0.0 | −1.1 | −2.2 | 1.1 | non-comp. |
| GYKI-53655 | NaN | −0.2 | NaN | >0.0 | −0.2 | – | non-comp. |
| GYKI-53784 | NaN | NaN | NaN | >0.0 | 0.0 | – | non-comp. |
| talampanel | NaN | NaN | NaN | NaN | 0.0 | – | non-comp. |

**Table S1**: Results for GluA2. All free energy values are provided in kcal/mol. NaN indicates that the simulation failed or the ligand was unstable in the binding pocket.

| Ligand | 8OP9 | 8RH7 | 8OQ7 | 9FRB | ΔΔG | Activity |
|---|---|---|---|---|---|---|
| (+)-CAMP | −8.02 | −9.44 | −7.87 | −7.85 | −1.57 | agonist |
| (R)-GABOB | −8.57 | −8.39 | −7.29 | −7.83 | −0.74 | agonist |
| (S)-GABOB | −9.46 | −8.78 | −6.66 | −7.84 | −1.62 | agonist |
| 5-Me-IAA | −8.56 | −5.65 | −5.44 | −6.32 | −2.24 | agonist |
| CACA | −7.30 | −10.57 | −4.74 | −5.80 | −4.77 | agonist |
| CGP36742 | 4.00 | 9.45 | −3.73 | −8.97 | 12.97 | antagonist |
| GABA | −8.74 | −12.19 | −7.55 | −6.66 | −4.64 | agonist |
| THIP | −4.19 | −6.96 | −8.03 | −5.46 | 1.07 | antagonist |
| TPMPA | −2.66 | 2.52 | −5.41 | −6.12 | 3.46 | antagonist |
| aza-THIP | −1.35 | 1.64 | −3.80 | −4.57 | 3.22 | antagonist |
| cis-3-ACPBPA | 5.32 | 23.55 | −3.57 | −5.49 | 10.81 | antagonist |
| isoguvacine | −6.31 | −2.38 | −5.74 | −5.65 | −0.57 | agonist |
| muscimol | −11.06 | −12.91 | −6.81 | −7.30 | −5.61 | agonist |
| piperidine-4-carboxylic acid (P4C, isonipecotic acid) | −4.49 | −5.26 | −5.40 | −4.57 | 0.14 | antagonist |
| piperidine-4-sulphonic acid (P4S) | −5.57 | −4.95 | −9.90 | −8.49 | 4.33 | antagonist |
| trans-3-ACPBPA | 4.68 | 12.07 | −3.12 | −8.56 | 13.24 | antagonist |

**Table S2**: Results for GABAAR ρ1. All free energy values are provided in kcal/mol.

| Ligand | $\Delta G_A$ | $\Delta G_I$ | $\Delta\Delta G$ | Activity |
|---|---|---|---|---|
| Cytisine | −11.22 | −9.58 | −1.64 | agonist |
| Nicotine | −11.84 | −10.76 | −1.08 | agonist |
| Epibatidine | −16.21 | −14.86 | −1.35 | agonist |
| AT-1001 | −18.74 | −19.63 | 0.89 | antagonist |
| SR-16584 | −11.48 | −14.46 | 2.98 | antagonist |
| 18-MC, Zolunicant | 19.31 | 8.10 | 11.21 | channel blocker |
| TC-2559 | −11.80 | −10.03 | −1.77 | agonist |
| Varenicline | −14.58 | −12.72 | −1.86 | agonist |
| PHA-543613 | −7.35 | −7.51 | 0.16 | antagonist |
| Carbachol | −11.04 | −10.71 | −0.33 | agonist |
| (R)-DHBE | −0.29 | −5.88 | 5.59 | antagonist |
| (S)-DHBE | 0.34 | −5.94 | 6.28 | |
| ABT-594, Tebanicline | −14.59 | −13.49 | −1.10 | agonist |
| NS-3861 | −16.08 | −15.58 | −0.50 | agonist |
| DMPP | −10.97 | −10.31 | −0.66 | agonist |

**Table S3**: Results for α3β4 nAChR. All free energy values are provided in kcal/mol.

| Ligand | $\Delta G_A$ | $\Delta G_I$ | $\Delta\Delta G$ | $E_{max}$ [%] | Ki [nM] |
|---|---|---|---|---|---|
| ALB125710 | −13.75 | −13.57 | −0.18 | 17 | 2.52 |
| ALB127594 | −12.41 | −12.39 | −0.03 | 16 | 1.1 |
| ALB133254 | −12.88 | −12.63 | −0.25 | 13 | 6.73 |
| ALB133255 | −10.32 | −11.42 | 1.09 | 14 | 33.7 |
| ALB133258 | −14.75 | −14.58 | −0.17 | 5 | 0.69 |
| ALB137637 | −9.80 | −10.47 | 0.66 | 10 | 108.6 |
| ALB148471 | −13.39 | −8.89 | −4.50 | 117 | 1.38 |
| Palonosetron | −13.28 | −15.38 | 2.10 | 0 | 0.246 |
| SMP100 | −12.40 | −11.13 | −1.27 | 41 | 2.26 |
| Serotonin | −12.34 | −7.99 | −4.36 | 100 | ~10 |

**Table S4**: Results for 5-HT3AR. All free energy values are provided in kcal/mol. The maximum response $E_{max}$ is given as a percentage of the reference ligand serotonin.

| Ligand | $\Delta G_A$ | $\Delta G_I$ | $\Delta\Delta G$ | Activity | $E_{max}$ [%] | EC50 [µM] | IC50 [µM] |
|---|---|---|---|---|---|---|---|
| compound 7 | −18.15 | −10.53 | −7.62 | agonist | 98 | 0.022 | >30 |
| compound 2 | −16.95 | −11.537 | −5.42 | agonist | 100 | 0.014 | >30 |
| compound 5 | −15.00 | −9.89 | −5.10 | agonist | 84 | 0.032 | >30 |
| compound 6 | −14.48 | −9.51 | −4.97 | agonist | 64 | 0.93 | >30 |
| compound 1a | −11.44 | −6.48 | −4.97 | agonist | 75 | 4.0 | >30 |
| compound 4a | −21.81 | −16.99 | −4.82 | agonist | 82 | 0.26 | >30 |
| compound 3 | −17.24 | −13.01 | −4.23 | agonist | 72 | 0.34 | >30 |
| compound 4b | −18.46 | −14.70 | −3.76 | (agonist) | – | >30 | >30 |
| compound 1b | −10.11 | −6.92 | −3.19 | (agonist) | – | >30 | >30 |
| compound 9b | −12.06 | −11.33 | −0.73 | (antagonist) | – | >30 | >30 |
| compound 10 | −16.38 | −16.52 | 0.14 | antagonist | – | >30 | 1.1 |
| compound 11 | −10.73 | −10.97 | 0.25 | antagonist | – | >30 | 11 |
| compound 13 | −10.24 | −11.33 | 1.08 | antagonist | – | >30 | 11 |
| compound 12 | −11.43 | −12.80 | 1.37 | antagonist | – | >30 | 12 |
| compound 9a | −12.98 | −16.04 | 3.06 | antagonist | – | >30 | 2.4 |
| compound 8 | −11.07 | −15.85 | 4.78 | antagonist | – | >30 | 4.4 |

**Table S5**: Results for TRPML1. All free energy values are provided in kcal/mol.

| Ligand | $\Delta G_A$ | $\Delta G_I$ | $\Delta\Delta G$ | Activity |
|---|---|---|---|---|
| Ebio2 | −18.16 | −15.03 | −3.13 | agonist |
| Ebio1 | −15.46 | −12.48 | −2.99 | agonist |
| RTG-S1 | −15.69 | −12.78 | −2.91 | agonist |
| QO-83 | −17.32 | −14.73 | −2.59 | agonist |
| CBD | −11.78 | −9.39 | −2.39 | agonist |
| HN37 | −18.53 | −16.22 | −2.31 | agonist |
| RTG | −12.82 | −10.62 | −2.20 | agonist |
| Ebio3 | −14.37 | −12.96 | −1.40 | antagonist |
| Ebio1-S1 | −12.96 | −12.66 | −0.31 | antagonist |

**Table S6**: Results for KCNQ2. All free energy values are provided in kcal/mol.
